# A tale of two gradients: Differences between the left and right hemispheres predict semantic cognition and visual reasoning

**DOI:** 10.1101/2021.02.23.432529

**Authors:** Tirso Rene del Jesus Gonzalez Alam, Brontë L. A. Mckeown, Zhiyao Gao, Boris Bernhardt, Reinder Vos de Wael, Daniel S. Margulies, Jonathan Smallwood, Elizabeth Jefferies

## Abstract

Decomposition of whole-brain functional connectivity patterns reveals a principal gradient that captures the separation of sensorimotor cortex from heteromodal regions in the default mode network (DMN); this gradient captures the systematic order of networks on the cortical surface. Functional homotopy is strongest in sensorimotor areas, and weakest in heteromodal cortices, suggesting there may be differences between the left and right hemispheres (LH/RH) in the principal gradient, especially towards its apex. This study characterised hemispheric differences in the position of large-scale cortical networks along the principal gradient, and their functional significance. We collected resting-state fMRI and semantic and non-verbal reasoning task performance in 175+ healthy volunteers. We then extracted the principal gradient of connectivity for each participant and tested which networks showed significant hemispheric differences in gradient value. We investigated the functional associations of these differences by regressing participants’ behavioural efficiency in tasks outside the scanner against their interhemispheric gradient difference for each network. LH showed a higher overall principal gradient value, consistent with its role in heteromodal semantic cognition. One frontotemporal control subnetwork was linked to individual differences in semantic cognition: when it was nearer heteromodal DMN on the principal gradient in LH, participants showed more efficient semantic retrieval. In contrast, when a dorsal attention subnetwork was closer to the heteromodal end of the principal gradient in RH, participants showed better visual reasoning. Lateralization of function may reflect differences in connectivity between control and heteromodal regions in LH, and attention and visual regions in RH.

## 1. Introduction

Contemporary accounts of brain organisation conceptualise cognition as reflecting interactions of large-scale networks of brain regions, organised in a systematic fashion along cortical gradients. These gradients capture similarities in connectivity patterns across disparate areas of the cortex (Bressler and Menon, 2010; Margulies et al., 2016; Medaglia et al., 2015; Paquola et al., 2018; Yeo et al., 2011). Cortical gradients provide a new tool for understanding patterns of hemispheric specialisation, since networks with lateralised connectivity will occupy different positions along these gradients in the left (LH) and right hemispheres (RH). This study exploits the potential of cortical gradients to uncover hemispheric differences in patterns of intrinsic connectivity, (i) by assessing the position of canonical networks in left and right hemisphere along gradients derived bilaterally, and (ii) by examining the functional significance of these hemispheric differences for semantic cognition and visual reasoning.

The principal gradient, which explains the most variance in whole-brain decompositions of intrinsic connectivity, captures the separation between sensory-motor cortex and heteromodal Default Mode Network (DMN) (Huntenburg et al., 2018; Margulies et al., 2016). In this way, it relates to previously described cortical hierarchies that extract progressively more complex or heteromodal information from sensory inputs, or that maintain more abstract goals for action, in lateral and medial temporal lobes, and lateral and medial prefrontal cortex (Badre, 2008; Badre and D’Esposito, 2007; Bajada et al., 2019, 2017; Fuster, 2001; Jackson et al., 2019, 2017; Koechlin et al., 2003; Petrides, 2005; Thiebaut De Schotten et al., 2017). The principal gradient goes beyond these observations to explain why similar hierarchies occur in multiple brain regions. The principal gradient is correlated with physical distance along the cortical surface from primary systems, with the DMN falling at a maximum distance from sensory and motor systems in multiple locations across the cortex. Since DMN is a highly distributed network, with multiple nodes located in distant brain regions, the functional transitions captured by the principal gradient are repeated across the cortex, and these are seen in both hemispheres. The principal gradient also captures the sequence of networks found along the cortical surface – from DMN, through frontoparietal control networks, to attention networks (Dorsal and Ventral, DAN and VAN) and finally primary somatomotor and visual networks. A recent study showed that when gradient decomposition is performed for the two hemispheres separately, both hemispheres contain a similar (but not identical) principal gradient (Liang et al., 2021). However, the functional relevance of these similarities and differences between the left and right hemisphere has not been established.

Patterns of intrinsic connectivity tend to be highly symmetrical, with the strongest time-series correlations seen between homotopic regions that occupy the same position in the two hemispheres (Jo et al., 2012). However, symmetrical patterns of connectivity are weaker within heteromodal networks towards the DMN apex of the gradient (Raemaekers et al., 2018). These increasing asymmetries are related to structural connectivity: primary cortices are connected across the hemispheres through fast fibres of the corpus callosum, while heteromodal cortices are connected by slower fibres that show less homotopic connectivity (Stark et al., 2008). A recent study using large-scale novel meta-analytic and voxel mirroring methods confirmed that areas with less similar connectivity across hemispheres are associated with heteromodal functions, such as memory, language and executive control (Mancuso et al., 2019). Moreover, higher-order networks, including DMN, frontoparietal network (FPN) and dorsal attention network (DAN), show the highest degrees of interhemispheric differences in intrinsic connectivity (Karolis et al., 2019; Wang et al., 2014). These lateralised patterns of connectivity have functional significance, giving rise to lateralised functions like language (Joliot et al., 2016; Knecht et al., 2000) and aspects of attention (Bartolomeo and Seidel Malkinson, 2019). For example, (Gotts et al., 2013) identified that a ‘segregation’ mode of lateralisation in LH (i.e., heightened intrinsic connectivity with other LH regions), conferred behavioural advantages in a verbal semantic task (vocabulary). In contrast, cross-hemisphere connections for RH were related to better visual reasoning (block design). Given that segregated connectivity is also associated with higher-order heteromodal networks, we would expect this LH-semantic pattern to involve lateralised connectivity at the heteromodal end of the gradient.

Previous studies have identified hemispheric differences in control networks, situated between DMN and sensory-motor cortex. In LH, the frontoparietal control network couples preferentially to DMN and language regions, while in RH, it shows stronger connectivity to attentional networks (Wang et al., 2014). These findings suggest that control networks might be critical for the emergence of lateralised cognition. In line with this view, the most lateralised regions of the semantic network are associated with controlled semantic retrieval, as opposed to conceptual representation (Gonzalez Alam et al., 2019). Furthermore, the clustering of connectivity patterns within the FPN across hemispheres reveals a bipartite organisation, with one subnetwork showing more intrinsic connectivity to DMN, whilst the other shows more connectivity to DAN (Dixon et al., 2018). These subnetworks may support the capacity of the FPN to couple efficiently with the DAN and DMN, depending on the task (Niendam et al., 2012; Spreng et al., 2013; Vincent et al., 2008; Wang et al., 2014). Collectively, these findings suggest that differences in network interactions between the hemispheres might be reflected in the location of control networks on the principal gradient, with LH control regions nearer to DMN, and RH control areas nearer to the sensory-motor end of the gradient. In line with this view, (Davey et al., 2016) suggested that left-lateralised semantic control processes reflect an interaction of conceptual knowledge, associated with DMN, and control processes that can promote the retrieval of currently-relevant aspects of knowledge, even when these are not dominant in long-term memory. Semantic cognition may be left lateralised because these DMN and control networks interact more strongly in the left hemisphere. In contrast, visual reasoning tasks are expected to involve an interaction of visual and control/attention networks (Hearne et al., 2017), without the strong engagement of memory processes in DMN. Gradient differences between LH and RH may promote these different patterns of network interaction such that networks involved in cognitive control can couple efficiently with both heteromodal DMN regions, towards the top of the principal gradient, and visual regions at the opposite end.

This study examines the organisation of the principal gradient across the left and right human cerebral hemispheres in participants who took part in a resting-state scan (N=253) and behavioural tasks on a separate session (N=175). First, we identify the position of 17 canonical networks on the principal gradient in the two hemispheres separately, using a widely-used decomposition of resting-state fMRI data (Yeo et al., 2011). This decomposition allows us to test the prediction that control networks in LH are situated closer to the heteromodal end of the principal gradient. Next, we use an individual differences approach to establish which gradient differences in connectivity relate to performance on tests of semantic cognition and visual reasoning. Finally, we characterise the response within LH and

RH portions of these lateralised networks to task manipulations of semantic control and working memory demands. Using similar methods, (Mckeown et al., 2020) found associations between individual differences in gradient values and patterns of spontaneous thought, suggesting that variation in gradient organisation is reflected in people’s experience. We build on these findings to relate hemispheric gradient differences to lateralised cognitive tasks.

## 2. Methods

### 2.1. Participants

Two hundred and seventy-seven healthy participants were recruited from the University of York. Written informed consent was obtained for all participants and the study was approved by the York Neuroimaging Centre Ethics Committee. Twenty-four participants were excluded from fMRI analyses; two due to technical issues during the neuroimaging data acquisition, one due to a data processing error and twenty-one for excessive movement during the scan (Power et al., 2014; mean framewise displacement > 0.3 mm and/or more than 15% of their data affected by motion), resulting in a final cohort of N = 253 (169 females, mean +/- SD age = 20.7 +/- 2.4 years). A subset of 175 of these participants also completed a semantic relatedness judgement task and Raven’s Progressive Matrices, in a separate session. While the current analysis of hemispheric gradient differences is novel, this data has been used in previous studies to examine the neural basis of memory and mind-wandering, including region-of-interest based connectivity analysis and cortical thickness investigations (Evans et al., 2020; Gonzalez Alam et al., 2018, 2019, 2021; Karapanagiotidis et al., 2017; Poerio et al., 2017; Sormaz et al., 2018; Turnbull et al., 2018; Vatansever et al., 2017; H. T. Wang et al., 2018; X. Wang et al., 2018).

### 2.2. Procedure

All participants underwent a 9-minute resting-state fMRI scan. During the scan, they were instructed to passively view a fixation cross and not to think of anything in particular.

Immediately following the scan, they completed a 25-item experience-sampling questionnaire while still in the scanner as part of separate studies; these have been reported in Karapanagiotidis et al. (2020) and Mckeown et al. (2020).

### 2.3. Materials

#### 2.3.1. Semantic Task

Participants performed semantic relatedness judgements that manipulated modality (words/pictures) and strength of association (weak/strong associates; see Figure 1). The task employed a three-alternative forced-choice design: participants matched a probe stimulus on the screen with one of three possible targets, and pressed buttons to indicate their choice. Each trial consisted of a centrally-presented probe preceded by a target and two unrelated distractors, which were targets in other trials. Trials started with a blank screen for 500ms. The three response options were subsequently presented at the bottom of the screen for 900ms (aligned horizontally, with the target in each location equally often). Finally, the probe was presented at the top of the screen. The probe and choices remained visible until the participant responded, or for a maximum of 3s. Both response time (RT) and accuracy were recorded, and an efficiency score was calculated for each participant in each condition by dividing response times by accuracy.

**Fig. 1.**
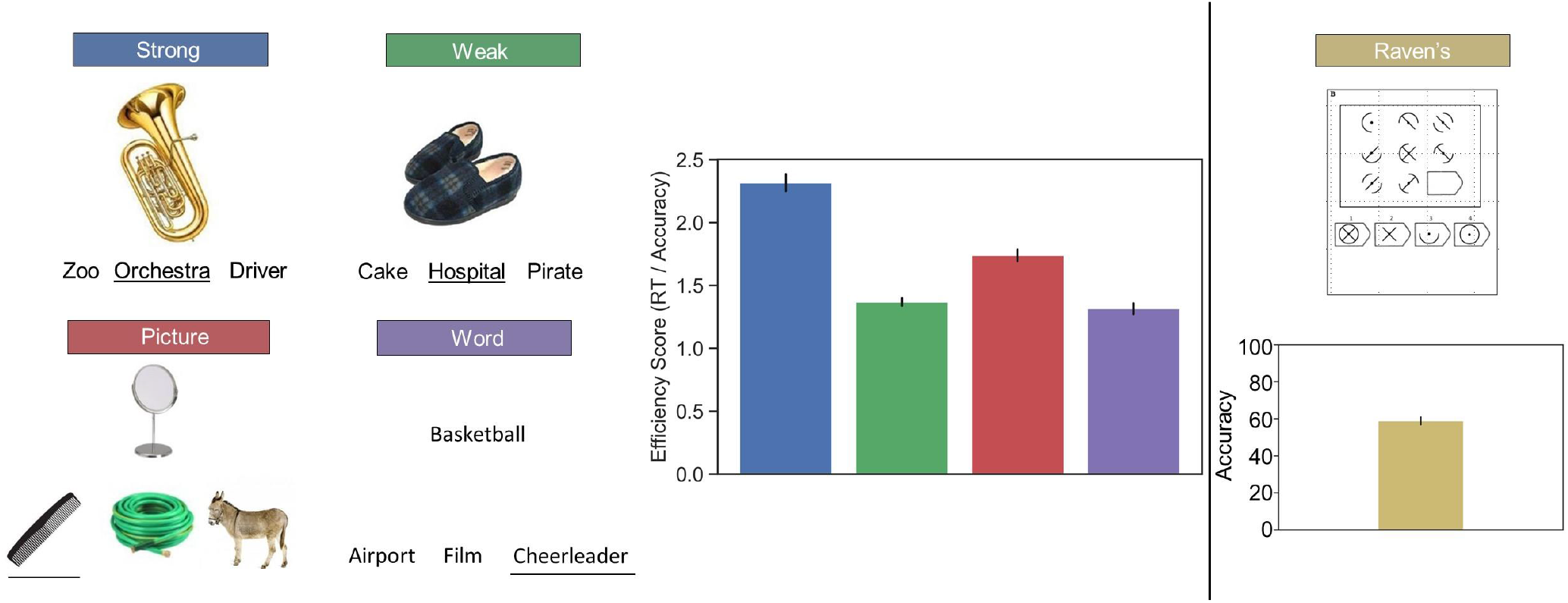
Illustration of the semantic (left panel) and non-semantic (right panel) tasks employed in this study. The bar plots are colour coded to match the colours of the boxes above the examples depicting each condition of the task (i.e., the blue bar in the left panel corresponds to the ‘strong’ condition). The error bars depict the 95% confidence interval

The stimuli employed in the tasks were selected from a larger set of words and photographs used in previous experiments (Davey et al., 2015; Krieger-Redwood et al., 2015). The pictures were coloured photographs collected from the internet and re-sized to fit the trial structure (200 pixels, 72 dpi). All the coloured pictures and words were rated for familiarity using 7-point Likert scales, and imageability (>500) from the MRC psycholinguistic database (Coltheart, 1981; Wilson, 1988). Lexical frequency for the words was obtained by the SUBTLEX-UK database (van Heuven et al., 2014) to allow matching on psycholinguistic properties. The strength of association between probe-target pairs was assessed using a 7-point Likert scale and differed significantly between conditions. There were no differences between strong and weak associations in word length, familiarity, imageability or lexical frequency. The order of trials within the blocks was randomized across subjects. The presentation of the blocks was interleaved.

##### 2.3.1.1. Semantic dimensionality reduction

Given that efficiency scores were correlated across the conditions of the task, we performed data-driven dimensionality reduction, which revealed a single semantic factor in the relatedness judgement task. PCA with varimax rotation yielded one single factor with Kaiser’s criterion above 1, explaining 75% of the variance. Each participant’s efficiency scores in the four tasks were therefore summarised using a single score reflecting the single factor loading, which was carried forward into regression analysis after z-scoring and imputing any outlier above +/- 2.5 with the mean.

#### 2.3.2. Non-Semantic Task: Raven’s Advanced Progressive Matrices

The Ravens Advanced Progressive Matrices (Raven et al., 1994) is a measure of non-verbal reasoning and required participants to identify meaningless visual patterns. The progressive matrices task included 36 questions, preceded by two practice trials. During the practice phase, participants were given feedback and task training, with no feedback for the reminder of the trials. For each problem, a set of 9 tiles (in a 3 × 3 design) were shown on the screen. All but one tile contained a pattern. At the bottom of the screen were 4 additional patterned tiles. Participants were required to select which tile would complete the pattern (see Figure 1). Participants were given 20 min to complete as many problems as they could, and the problems got progressively more difficult.

### 2.3. Neuroimaging

The MRI data acquisition and pre-processing steps reported in this paper are identical to the steps reported in Karapanagiotidis et al. (2020), and the dimension reduction steps are identical to the ones reported in Mckeown et al. (2020), as reproduced in the sections below.

#### 2.3.1. MRI Data Acquisition

MRI data were acquired on a GE 3 T Signa Excite HDx MRI scanner, equipped with an eight-channel phased array head coil at York Neuroimaging Centre, University of York. For each participant, we acquired a sagittal isotropic 3D fast spoiled gradient-recalled echo T1-weighted structural scan (TR = 7.8 ms, TE = minimum full, flip angle = 20°, matrix = 256×256, voxel size = 1.13 x 1.13 x 1 mm^3^, FOV = 289 x 289 mm^2^). Resting-state fMRI data based on blood oxygen level-dependent contrast images with fat saturation were acquired using a gradient single-shot echo-planar imaging sequence (TE = minimum full (≈19 ms), flip angle = 90°, matrix = 64×64, FOV = 192 x 192 mm^2^, voxel size = 3×3×3 mm^3^, TR = 3000 ms, 60 axial slices with no gap and slice thickness of 3mm). Scan duration was 9 minutes which allowed us to collect 180 whole-brain volumes.

#### 2.5.2. MRI data pre-processing

fMRI data pre-processing was performed using SPM12 (http://www.fil.ion.ucl.ac.uk/spm) and the CONN toolbox (v.18b) (https://www.nitrc.org/projects/conn) (Whitfield-Gabrieli and Nieto-Castanon, 2012) implemented in Matlab (R2018a) (https://uk.mathworks.com/products/matlab). Pre-processing steps followed CONN’s default pipeline and included motion estimation and correction by volume realignment using a six-parameter rigid body transformation, slice-time correction, and simultaneous grey matter (GM), white matter (WM) and cerebrospinal fluid (CSF) segmentation and normalisation to MNI152 stereotactic space (2 mm isotropic) of both functional and structural data. Following pre-processing, the following potential confounders were statistically controlled for: 6 motion parameters calculated at the previous step and their 1st and 2nd order derivatives, volumes with excessive movement (motion greater than 0.5 mm and global signal changes larger than z = 3), linear drifts, and five principal components of the signal from WM and CSF (CompCor approach; Behzadi et al., 2007). Finally, data were band-pass filtered between 0.01 and 0.1 Hz. No global signal regression was performed.

#### 2.3.2. Gradient analysis

We obtained each participant’s gradient values for the first principal gradient (Margulies et al., 2016) following the methods described in Mckeown et al. (2020). Following pre-processing, the functional time-series from 400 ROIs based on the Schaefer parcellation (Schaefer et al., 2018; Yeo et al., 2011) were extracted for each individual. A connectivity matrix was then calculated using Pearson correlation resulting in a 400×400 connectivity matrix for each participant. These individual connectivity matrices were then averaged to calculate a group-averaged connectivity matrix. The BrainSpace Toolbox (Vos de Wael et al., 2020) was used to extract ten group-level gradients from the group-averaged connectivity matrix (dimension reduction technique = diffusion embedding, kernel = normalized angle, sparsity = 0.9). This study was primarily focussed on the first gradient, which has well-described functional associations relevant to previous lateralisation findings; however, we extracted ten gradients to maximize the degree of fit between the group-averaged gradients and the individual-level first gradient (Supplementary Table 1 in Mckeown et al., 2020, shows the average degree of fit for gradient one when extracting ten gradients compared to three). The variance explained by each group-averaged gradient is provided in Mckeown et al. (2020) Supplementary Figure 1.

The group-level gradient solutions were aligned using Procrustes rotation to a subsample of the HCP dataset (n = 217, 122 women, mean +/- SD age = 28.5 +/- 3.7 y; for full details of subject selection see (Vos De Wael et al., 2018) within the BrainSpace toolbox (Vos de Wael et al., 2020). The first three group-averaged gradients, with and without alignment to the HCP data are shown in Supplementary Figure 2 of Mckeown et al. (2020). To demonstrate the benefits of this alignment step, we calculated the similarity using Spearman Rank correlation between the first five aligned and unaligned group-level gradients to the first five gradients reported in Margulies et al. (2016) which were calculated using 820 participants over an hour resting-state scan. Alignment improved the stability of the group-level gradient templates by maximising the comparability of the solutions to those from the existing literature (i.e., Margulies et al., 2016; see Supplementary Table 2 in Mckeown et al., 2020).

Using identical parameters, individual-level gradients were then calculated for each individual using their 400×400 connectivity matrix. These individual-level gradient maps were aligned to the group-level gradient maps using Procrustes rotation to improve the comparison between the group-level gradients and individual-level gradients (N iterations = 10). This analysis resulted in ten group-level gradients and ten individual-level gradients for each participant explaining maximal whole-brain connectivity variance in descending order. As stated above, this report focuses on the principal gradient (with supplementary analyses for Gradient 2). To demonstrate the variability of individual-level gradients, Supplementary Figure 3 in Mckeown et al. (2020) shows the highest, lowest, and median similarity gradient maps for the principal gradient.

#### 2.3.3. Hemispheric Difference Analysis

As a first step for our analysis of interest, we obtained group averages of the principal gradient for each of the 400 parcels per participant (top row of Figure 3). Since these parcels do not necessarily share homotopes across hemispheres, for the hemispheric difference analyses we summarised these values by averaging, for each participant, the parcels corresponding to each of the 17 networks described by Yeo et al. (2011). We will refer to these two levels of analyses as ‘parcel level’ and ‘network level’ respectively.

Next, we examined hemispheric differences across the 17 Yeo Network parcellation. We normalised each network’s average principal gradient value within each participant using a minimum-maximum normalisation (0-100) at the parcel level, such that networks toward the lower end of the principal gradient have values closer to 0, and networks towards DMN have values close to 100 (the middle row of Figure 3 shows the group average per network; the organisation of these networks are depicted in the bottom row of Figure 3). We tested for hemispheric differences in the global gradient value by averaging all gradient values across the 17 Yeo networks within each hemisphere separately for each participant in the sample and comparing these means using a paired t-test (LH vs RH).

Our next step involved subtracting the average of each RH network from its homotope in the LH, per participant (we z-scored the results to produce a group difference map highlighting the networks with the most extreme differences shown in Figure 4). We then performed a two-way repeated measures ANOVA, using Hemisphere and Network as between-subject factors, to test for hemispheric differences at the network level. Having obtained significant main effects and an interaction, we conducted post-hoc non-parametric permutation testing with 5000 bootstrapped samples to compute the probability of obtaining a difference of gradient means across hemispheres as extreme as that empirically observed for each network by chance (Figure 5). The non-parametric p-values of these post-hoc tests were Bonferroni-corrected at an alpha=0.05 for 17 multiple comparisons to guard against Type 1 errors. We only included those networks that showed significant hemispheric differences in the subsequent analyses.

#### 2.3.4. Behavioural Regressions

In order to examine whether hemispheric differences on the principal gradient across networks had behavioural consequences, we performed regression analyses relating participants’ performance outside the scanner on semantic and visual reasoning tasks to the difference in principal gradient values across the hemispheres for each significant network. We entered each participant’s semantic factor loading and their z-scored performance on the Raven’s task as Explanatory Variables (EVs) into an Ordinary Least Squares (OLS) regression using hemispheric difference scores on the principal gradient as the dependent variable.

### 2.4. Parametric manipulations of semantic control and working memory load

In our final analysis, we examined parametric maps reflecting the effect of control demands in semantic judgements and in verbal working memory task in N=27 participants (a re-analysis of data in (Gao et al., 2020). This analysis established whether the networks that showed an association with individual differences in cognition differed in terms of their response to semantic and non-semantic control demands in the LH and RH. We would expect a lateralised response to semantic but not non-semantic control demands for networks in which hemispheric differences in principal gradient values predicted individual differences in semantic cognition – i.e. a stronger response to semantic control than to other task demands in LH but not RH. We would not expect this pattern for networks showing a lateralised response to visual reasoning. These participants did not take part in the resting-state fMRI session.

In the on-line semantic task in fMRI, participants had to judge whether pairs of words were semantically related or unrelated. Each trial consisted of a word followed by a fixation cross and then a second word; then a blank screen signalled the decision period which marked the end of the trial. The stimuli had varying degrees of thematic relatedness depending on their frequency of co-occurrence (no taxonomically related stimuli were included). The degree of relatedness was quantified using distributed representations of word meanings obtained from the word2vec neural network, trained on the 100 billion-word Google News dataset (Mikolov et al., 2013). The stimulus set was manipulated so there was a continuum of relatedness, from ‘not related at all’ to ‘strongly related’. Since the degree of relatedness was continuous, there were no clear ‘correct’ or ‘incorrect’ answers; instead, the trials were sorted according to whether participants judged each trial to be related or unrelated. Difficulty was then estimated for related and unrelated trials separately by binning the stimuli into 5 categories according to their word2vec score. For trials judged to be related, a lower word2vec score was associated with increased difficulty (since establishing a semantic link for less strongly related items is harder); conversely, for trials judged to be unrelated, a higher word2vec score increased difficulty (since rejecting a relationship between associated words is harder).

Non-semantic control demands were manipulated in a verbal working memory task, where the parametric manipulation consisted of the number of items participants had to maintain in memory. This task had a similar structure and method of presentation to the semantic task; each trial began with a letter string (3 to 7 letters) presented at the centre of the screen, followed by a fixation. Participants were asked to remember these letters. Next, two letters were shown on the screen and participants judged whether both of them had been presented in the letter string. The working memory load was manipulated by varying the number of letters memorised in each trial; there were five levels of load from 3 to 7 letters.

The univariate analysis of these parametric manipulations yielded effect maps for semantic control and working memory demands, which we employed in an ROI analysis to examine which networks showed significant task differences across hemispheres. For the participant level analysis, we binarised the Yeo network maps and used them as ROI masks to extract the percent signal change value for each of the 27 participants separately in LH and RH for each condition of the task (related, unrelated and working memory) using the featquery tool in FSL 6. We entered these values per participant into separate repeated measures ANOVAs for each network using ‘Hemisphere’ and ‘Condition’ as within-subjects factors.

### 2.5. Supplementary Analysis of the Second Gradient

While our main focus is on the principal gradient, we provide a supplementary analysis of the second gradient as described in Margulies et al. (2016), which captures the difference in connectivity between visual and motor networks. We show the means per hemisphere for gradient 2, along with group means for each of the 400 parcels from Schaefer et al. (2018), and the 17 networks from Yeo et al. (2011). We characterise hemispheric differences per network for this gradient. Lastly, we provide bootstrapping analyses of the LH versus RH network differences and establish which networks survive correction for multiple comparisons (see Supplementary Analysis: Gradient 2). All of these analyses follow the methods described above.

## 3. Results

### 3.1. Gradient values for the Schaeffer parcellation in left and right hemispheres

Our first analysis step revealed a global mean difference on the principal gradient (see Figure 2), with higher values in the left hemisphere (paired samples t-test: t(252) = 18.38, p < .001). Participants’ LH and RH mean gradient values were very highly correlated (Pearson’s r = 0.9, p < .001), despite this global difference.

**Fig. 2.**
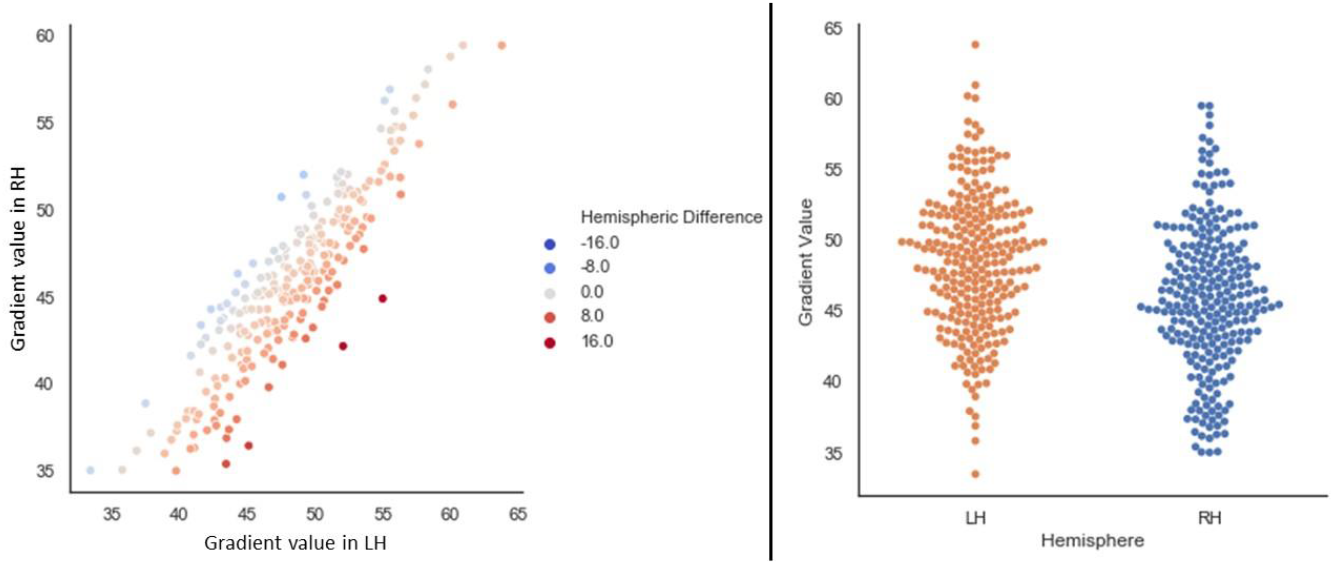
The left panel depicts a linear relationship in our sample’s mean left and right hemisphere values on the principal gradient. The ‘Hemispheric Difference’ legend of the scatterplot depicts the result of subtracting the LH – RH gradient loadings for the whole hemisphere per participant. Positive values reflect closer proximity to the heteromodal end of the gradient in LH. Negative values reflect closer proximity to the heteromodal end of the gradient in RH. The right panel depicts the distributions of mean global hemispheric values per participant in our sample. In both plots, each dot represents one participant. The scale on both plots indicates values on the principal gradient, which were re-scaled to range from 0 to 100

As expected, given our gradient alignment methods, the gradient decomposition of our 253-participant sample showed a principal gradient very similar to the one reported by Margulies et al. (2016), both at the parcel level (Figure 3, top row) and at the network level (Figure 3, middle row); see also Mckeown et al. (2020).

**Fig. 3.**
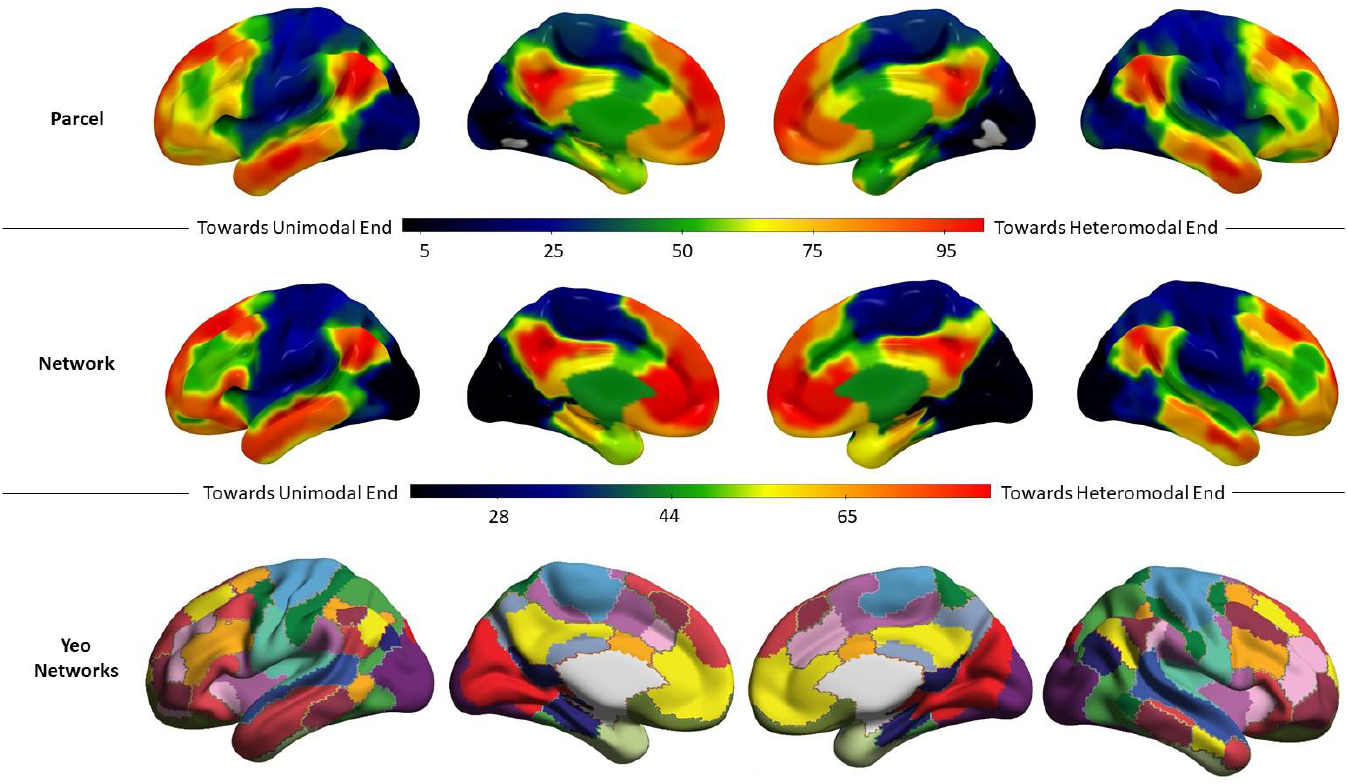
Top row: Group mean principal gradient value for each parcel in Schaeffer’s 400-parcel solution for our sample of 253 participants. Middle row: Group mean principal gradient value for each network in Yeo’s 17-network solution for our sample of 253 participants. Gradient units are arbitrary and have been normalised on a 0-100 minmax scale. Bottom row: 17 network parcellation by Yeo et al. (2011; the colour code followed in this figure replicates that of Buckner et al., 2011)

### 3.2. Hemispheric difference analysis at the network level

In order to compare the principal gradient loadings of the regions captured in the 400-region parcellation across the cerebral hemispheres, we averaged all parcels that fell within each network in the left and right hemispheres separately, and then performed a subtraction (LH - RH) and z-scored the resulting differences. The results can be seen in Figure 4. The principal gradient loadings in warm colours are nearer the heteromodal apex in the LH compared to RH, and the cool colours represent principal gradient loadings that are nearer the heteromodal apex in RH compared to LH. The value of these LH - RH network gradient differences was highly correlated with the principal gradient at the group level (r = 0.93, p < .0001), consistent with the expectation that heteromodal cortex shows more divergent connectivity across the hemispheres than unimodal cortex.

**Fig. 4.**
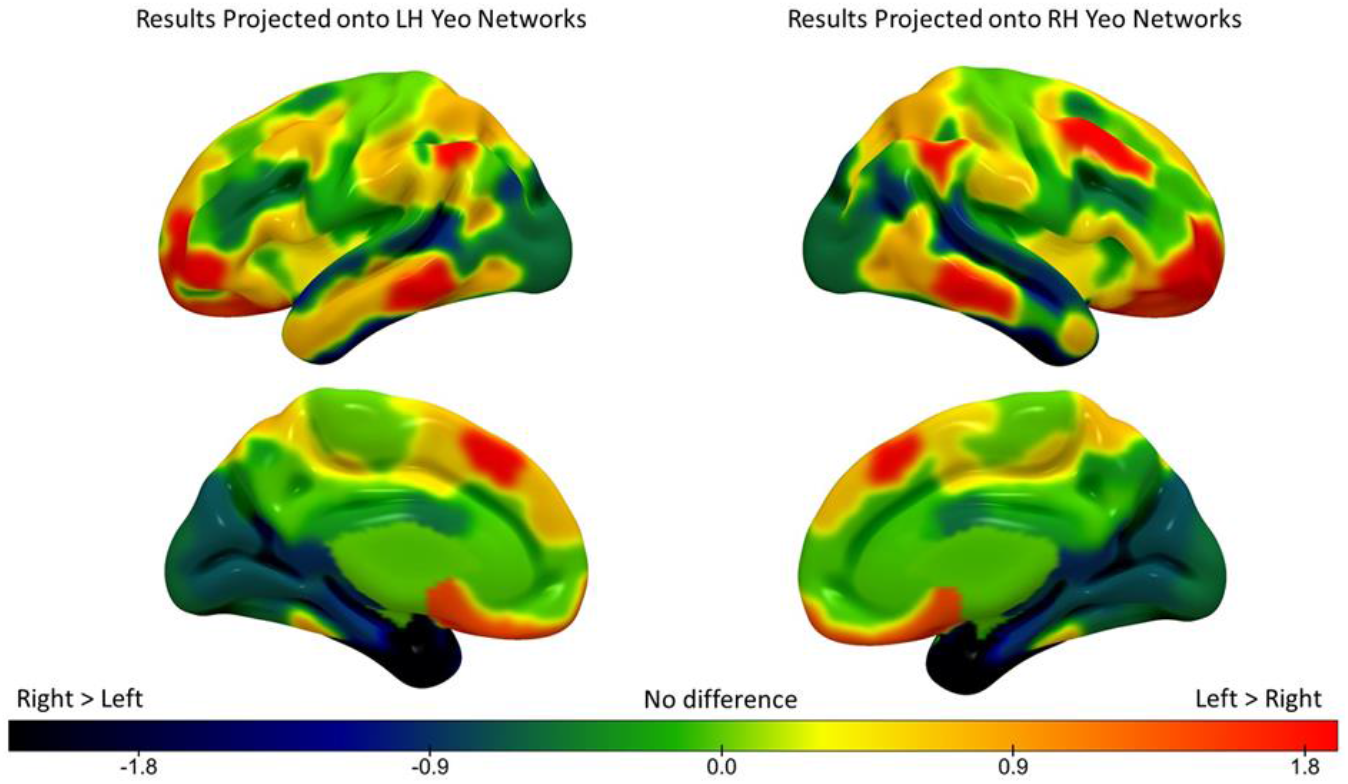
Hemispheric differences in principal gradient values across the 17 Yeo networks (z-scored). The warm colours represent principal gradient loadings that are nearer the heteromodal apex in LH compared to RH. Cool colours represent principal gradient loadings that are nearer the heteromodal apex in RH compared to LH

We performed a repeated-measures ANOVA to formally test for differences in principal gradient loadings at the network level (2 hemispheres by 17 networks), controlling for global hemispheric differences in gradient values by entering each participant’s global LH – RH difference value as a covariate of no interest. The results of this ANOVA revealed significant main effects of hemisphere (F(1,251) = 538.82, p < .0001, ηp2 = .68), and network (F(8.48, 2127.52) = 902.44, p < .0001, ηp2 = .78), as well as a significant hemisphere by network interaction (F(10.62, 2664.95) = 18.61, p < .0001, ηp2 = .07; all values with Greenhouse-Geisser correction to account for violations of the sphericity assumption). Subsequent post-hoc tests comparing LH and RH for each network (using permutation testing with 5,000 simulations to establish significance; Bonferroni-corrected for 17 comparisons) revealed that these hemispheric differences were robust for seven networks: DMN-B, Control B, Limbic B, Limbic A, DAN-A, DAN-B, and VAN-A (Figure 5). Only these seven networks were carried forward for further analyses. Supplementary Figure 1 shows the distribution of gradient values for these seven networks.

**Fig. 5.**
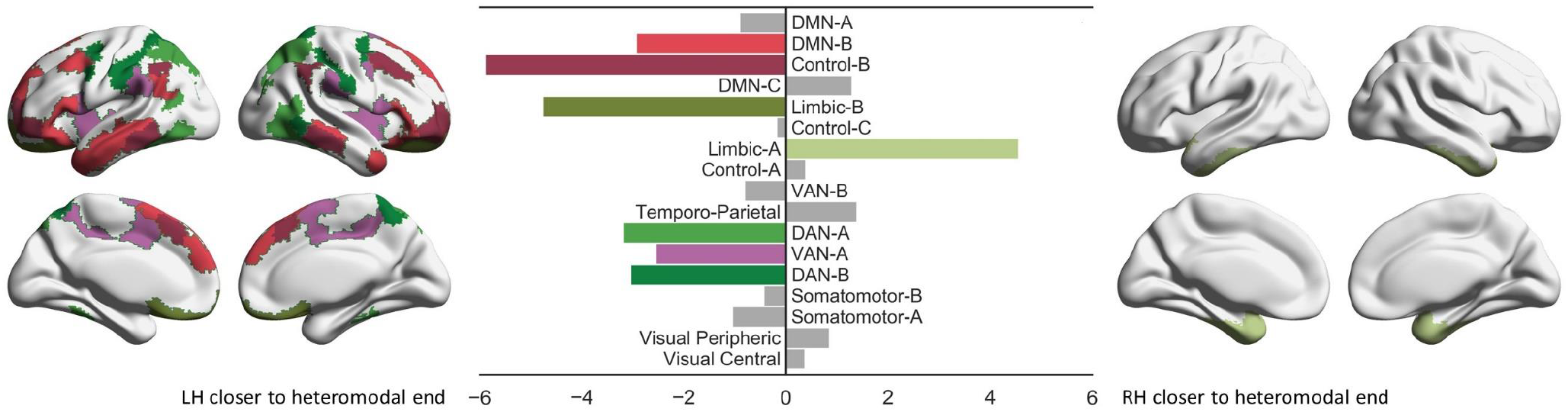
Results of permutation testing of LH versus RH positions on the principal gradient for each network (5,000 simulations, Bonferroni-corrected alpha for 17 comparisons). The size of each bar reflects the normalized (0-100) empirically observed mean difference across the hemispheres for each network. Coloured bars denote networks that showed significant differences (see Table 1 for exact p-values) and are colour-coded to indicate the position of each network in the brain. The brain map on the left side of the plot shows networks that were closer to the heteromodal end of the gradient in LH, while the brain map on the right side of the plot shows one network that was closer to the heteromodal end of the gradient in RH

**Table 1.** Normalised (on a scale of 0-100) means across hemispheres for each Yeo network. Larger values reflect greater proximity to the heteromodal end of the principal gradient. The p values indicate the results of pairwise bootstrapped permutation testing of LH vs RH principal gradient means for each network (5,000 simulations). Note: *** = significant at p < 0.0002 with Bonferroni-correction for 17 comparisons.

| **Network** | **LH Mean (SD)** | **RH Mean (SD)** | **Corrected p value** |
| --- | --- | --- | --- |
| Default A | 82.76 (5.84) | 81.85 (5.5) | > 0.1 |
| Default B | 77.62 (6.2) | 74.69 (6.88) | .000*** |
| Control B | 73.29 (7.51) | 67.39 (7.44) | .000*** |
| Default C | 64.75 (9.98) | 66.05 (10.4) | > 0.1 |
| Limbic B | 67.51 (10.38) | 62.74 (9.54) | .000*** |
| Control C | 59.6 (12.14) | 59.41 (11.38) | > 0.1 |
| Limbic A | 54.66 (9.51) | 59.22 (8.53) | .000*** |
| Control A | 52.37 (7.54) | 52.76 (7.59) | > 0.1 |
| Salience / Ventral Attention B | 50.67 (9.18) | 49.85 (9.18) | > 0.1 |
| Temporo-Parietal | 48.74 (11.07) | 50.14 (10.37) | > 0.1 |
| Dorsal Attention A | 39.03 (8.61) | 35.84 (8.35) | .000*** |
| Salience / Ventral Attention A | 35.94 (7.37) | 33.38 (7.29) | .000*** |
| Dorsal Attention B | 34.11 (7.57) | 31.07 (7.3) | .000*** |
| Somatomotor B | 29.47 (7.33) | 29.03 (7.38) | > 0.1 |
| Somatomotor A | 29.56 (7.29) | 28.51 (7.47) | > 0.1 |
| Visual Peripheric | 18.84 (8.96) | 19.69 (8.73) | > 0.1 |
| Visual Central | 19.11 (6.57) | 19.48 (7.04) | > 0.1 |

### 3.3. Behavioural Regressions

We next tested whether the degree of difference in principal gradient loadings across hemispheres for each Yeo network was associated with the efficiency with which participants performed semantic and visual reasoning tasks outside the scanner. We defined regression models using the empirically observed mean hemispheric difference in gradient scores (LH - RH) for each network as the dependent variable, and the efficiency of semantic decisions and accuracy on Raven’s matrices as two explanatory variables per participant. There was a significant association between task performance and hemispheric gradient differences for two out of seven networks (only networks showing a significant difference on the principal gradient in the analysis above were included). Hemispheric differences in gradient values for Control-B showed a positive association with overall semantic performance (β = .19, p = .007), and no relationship with visual reasoning (β = .01, p > .1). DAN-B showed a negative association between LH – RH gradient loadings and visual reasoning (β = −.15, p = .022) and no relationship with semantic performance (β = .02, p > .1) (see Figure 6). Participants whose Control-B network was closer to the heteromodal DMN end of the principal gradient in LH compared with RH showed more efficient semantic retrieval; in contrast when the DAN-B network was closer to the heteromodal end of the principal gradient in RH compared with LH, participants showed better visual reasoning on a matrices task. An additional analysis comparing subtasks of the semantic battery observed no significant association between gradient hemispheric differences and the effects of modality of presentation (pictures versus words) or strength of association (weak versus strong associations), with all p values > 0.1 (see supplementary materials, Table S1).

**Fig. 6.**
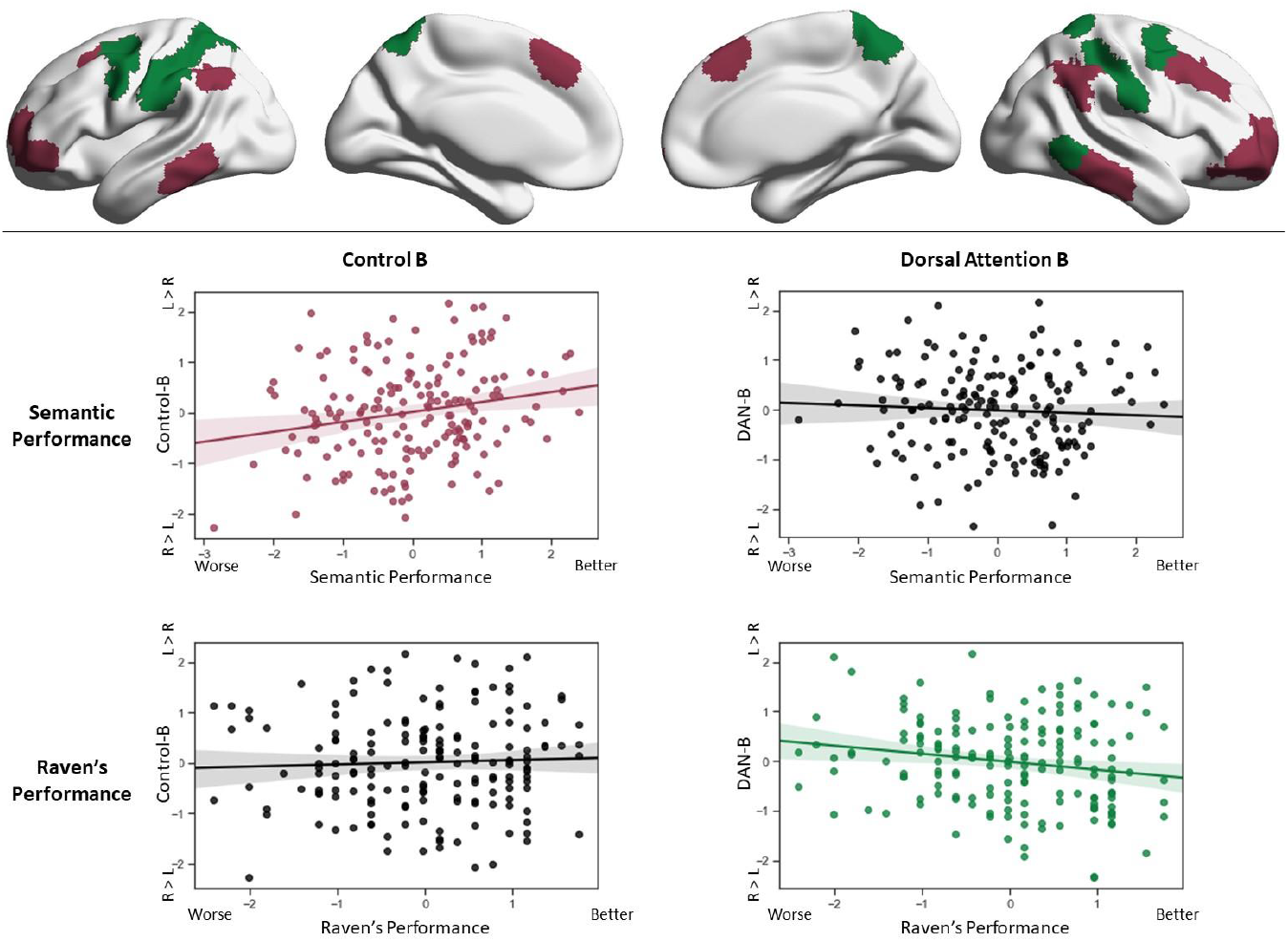
Scatterplots showing the relationship between hemispheric difference scores on the principal gradient and efficiency on semantic decisions (middle row) and accuracy on a visual reasoning task (Raven’s progressive matrices; bottom row). Only networks with significant results are shown (Control B on the left-hand side; Dorsal Attention B on the right-hand side). The scatterplots in colour denote significant effects in the regression model and have been colour coded to the networks driving the effect, shown in the top row

Given that hemispheric differences in principal gradient values predicted individual differences in semantic cognition, we next asked if Control-B and DMN networks are closer on the principal gradient of connectivity in LH compared with RH. This finding would be consistent with the greater coupling of these networks in states of controlled semantic retrieval reported by Davey et al. (2016). Yet the sensitivity of the principal gradient to this pattern of functionally-relevant network similarity in the LH is not yet established, since both Control-B and DMN-B (the adjacent network on the principal gradient) showed higher gradient values for LH compared with RH in the analysis in Figure 5.

We computed the distance on the principal gradient between Control-B and the two networks that were closer to the heteromodal apex of the principal gradient (DMN-A and DMN-B) for each hemisphere separately, and compared these distances across hemispheres using paired t-tests (all p values reported are corrected for multiple comparisons). There was a significantly smaller difference in principal gradient values between Control-B and DMN-B in LH compared with RH (mean difference in LH = 4.33, SD = 7.4; and in RH = 7.29, SD = 7.93; t(174) = 6.52, p < .001). There was a similarly smaller difference in principal gradient values between Control-B and DMN-A in LH compared with RH (mean difference in LH = 9.47, SD = 7.77; and in RH = 14.45, SD = 7.06; t(174) = 13.07, p < .001). This confirms that Control-B is closer to DMN along the principal gradient.

We repeated this analysis to establish if DAN-B has greater proximity to sensorimotor networks in RH compared with LH. There were significantly smaller gradient distances in RH compared with LH for all four relevant network comparisons: (i) visual central: LH = 15.01, SD = 8.7; RH = 11.6, SD = 9.09; t(174) = 7.96, p < .001); (ii) visual peripheral: LH = 15.28, SD = 11.33; RH = 11.37, SD = 11.47; t(174) = 8.76, p < .001), (iii) somatomotor-A (LH = 4.55, SD = 5.93; RH = 2.55, SD = 5.26; t(174) = 7.95, p < .001) and (iv) somatomotor-B (LH = 4.64, SD = 5.12; RH = 2.04, SD = 4.88; t(174) = 9.75, p < .001). DAN-B was closer to all sensorimotor networks in the RH compared with the LH. These results can be seen in Figure 7.

**Fig. 7.**
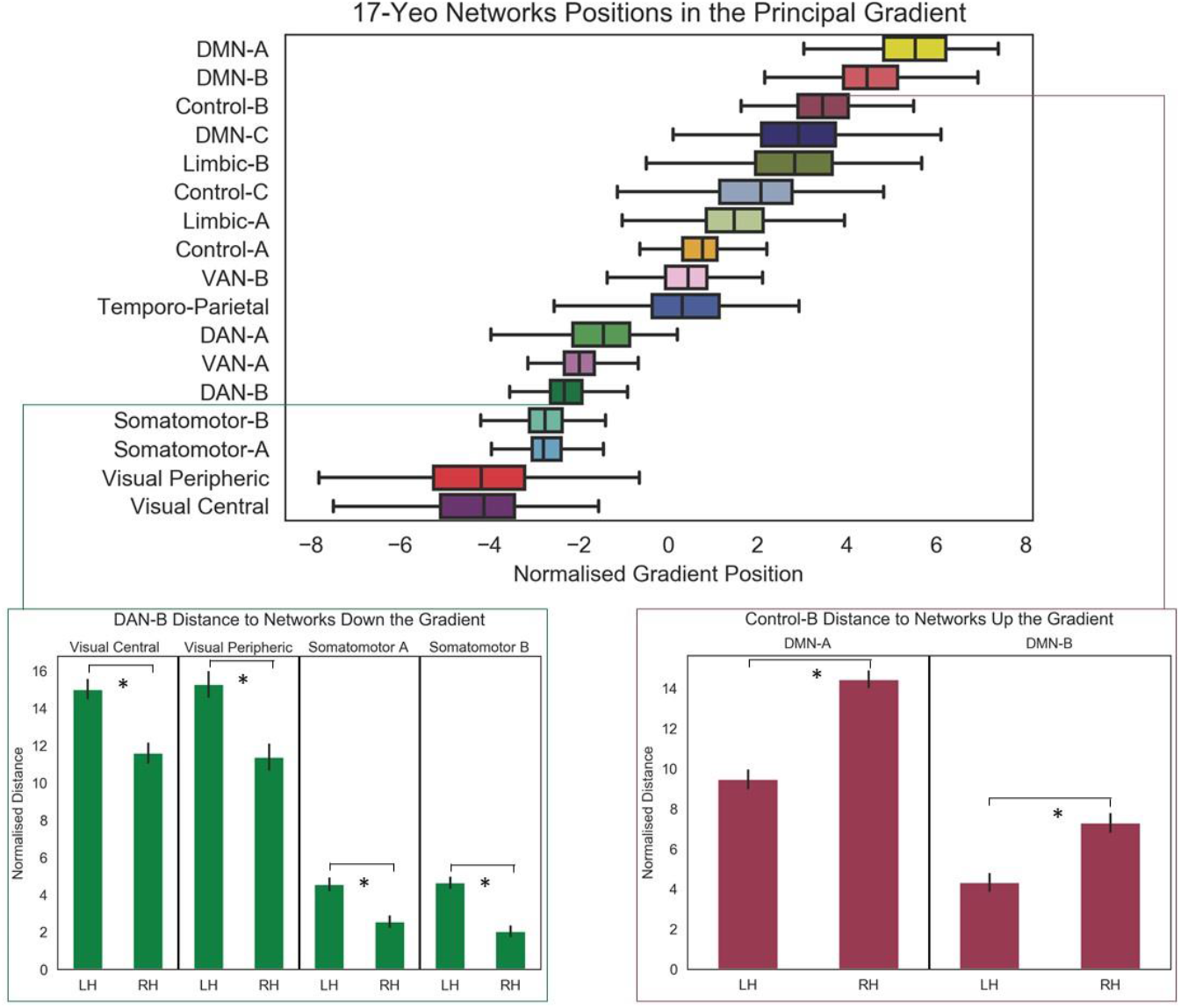
Differences of network positions in the principal gradient between pairs of networks that are relevant for lateralised cognitive processes. Note: * = p < .001 (corrected for multiple comparisons)

### 3.4. Parametric Effects of Control Demands

The individual differences analysis found that when Control-B is closer to the DMN-end of the principal gradient in LH versus RH, participants have more efficient semantic retrieval. In contrast, when DAN-B is closer to the unimodal end of the principal gradient in RH, participants show better visual reasoning on a progressive matrices task. These findings predict a hemispheric dissociation between networks in the effects of control demands across domains (i.e. in effects of semantic control demands and non-semantic difficulty – even within the verbal domain). We tested this prediction by examining the effects of parametric manipulations of semantic control demands (strength of association) and verbal working memory load on activation within the LH and RH components of control-B and DAN-B networks. An omnibus ANOVA examining the factors of hemisphere (LH vs. RH), task difficulty (related semantic, unrelated semantic and working memory) and network (Control-B vs. DAN-B) showed a three-way interaction between these factors (F(2,54) = 6.71, p = 0.003).

Separate repeated-measures ANOVAs for control-B and DMN-B found distinct patterns. Control B showed a significant interaction between hemisphere and condition, reflecting larger effects of control demands in the semantic task relative to working memory (F(1.34,36.07) = 7.72, p = .005, ηp2 = .22). Post-hoc tests revealed a greater response to difficulty for semantic decisions in LH versus RH (p < .001); in contrast, there were no hemispheric differences in the effect of WM load. There was no task by hemisphere interaction for DAN-B (F < 1). There were also no main effects of task in either network, but there was more activation in LH overall, likely reflecting the verbal nature of the tasks. Full ANOVA results are reported in Table 2. These results confirm that the left-lateralised components of Control-B show a specific response to semantic control demands, but not to working memory load. In contrast, DAN-B shows an equivalent response to the two forms of verbal control across hemispheres. These effects are shown in Figure 8.

**Table 2.**
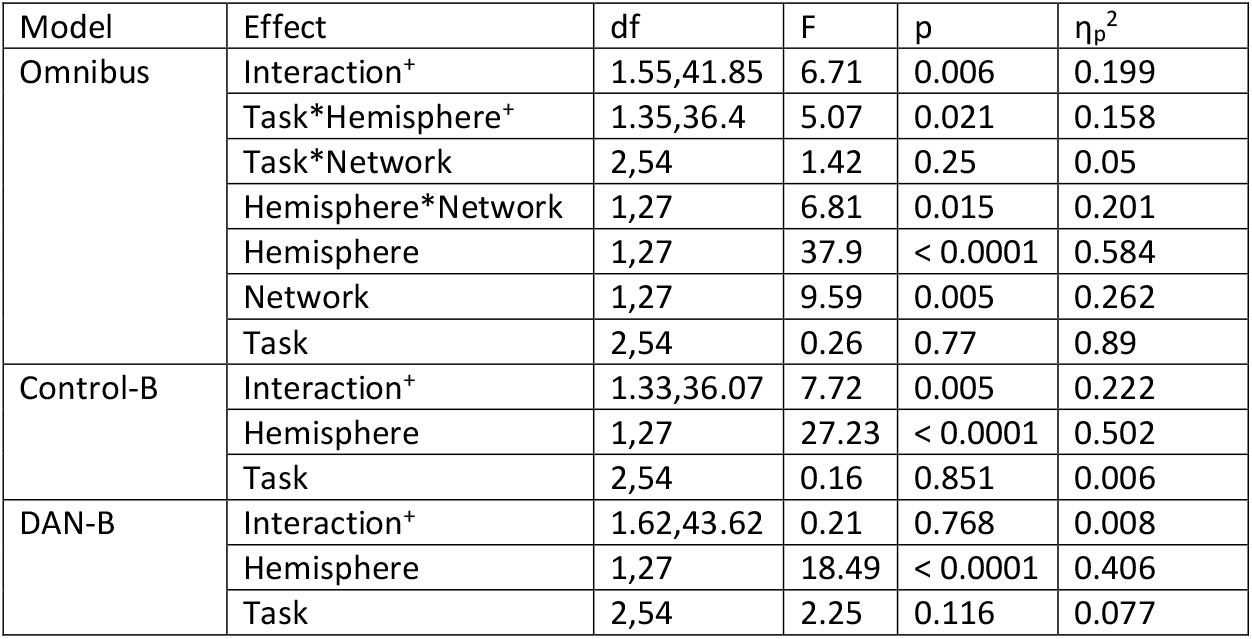
Results of the ANOVAs on the parametric difficulty effect maps. Note. The effects marked with + were subjected to Greenhouse-Geisser corrected since our data violated the assumption of sphericity (Mauchley’s test of sphericity p < .05 in both cases).

**Fig. 8.**
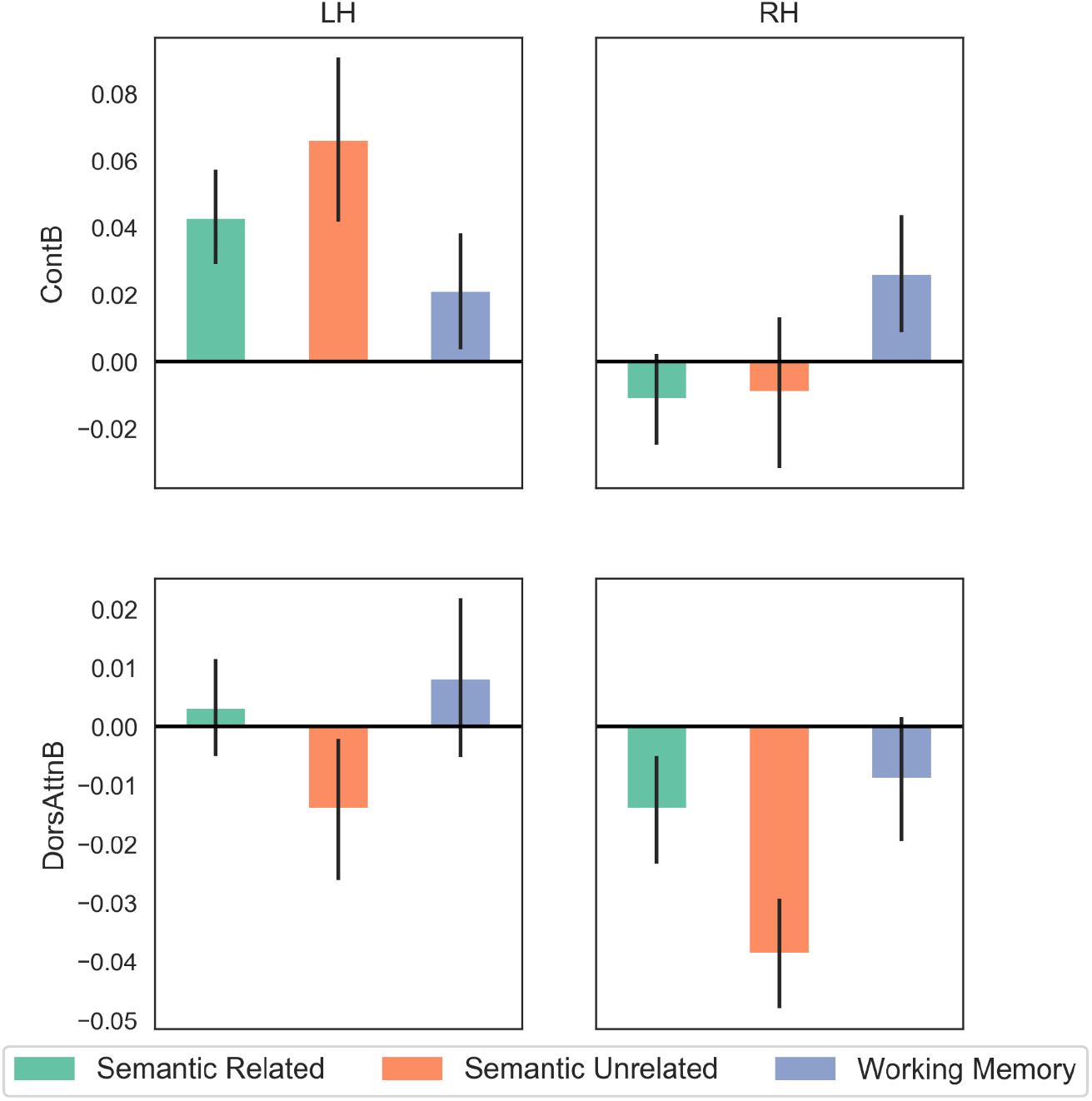
Parametric effects of difficulty in semantic and non-semantic tasks for 27 participants in Control-B and Dorsal Attention-B Yeo networks, split by hemisphere (the error bars depict the standard error of the mean), with effects expressed in percentage signal change

## 4. Discussion

This study investigates the lateralisation of function along the principal gradient – a key topographical component of large-scale intrinsic connectivity patterns capturing the separation of unimodal and heteromodal cortex (Margulies et al., 2016). We show that intrinsic connectivity patterns in the two hemispheres are situated at different points along the principal gradient: overall, LH parcels are closer to the heteromodal end of the principal gradient than RH parcels, consistent with the role of this hemisphere in key heteromodal functions, such as semantic cognition and language. This pattern was observed in many canonical heteromodal networks derived from a whole-brain parcellation of resting-state data (Yeo et al., 2011), including control, default, dorsal and ventral attention networks, but we did not observe a gradient difference between the hemispheres in sensorimotor networks. This pattern was also inverted for Limbic-A, centred on the ventral anterior temporal lobe (ATL), since for this network, the RH was closer to the heteromodal end of the principal gradient. In addition, individual differences in the relative gradient positions of networks across the hemispheres were found to have functional consequences for two cognitive processes with opposing patterns of lateralisation, semantic cognition and visual reasoning. Participants whose Control-B network was closer to the heteromodal DMN end of the principal gradient in LH compared with RH showed more efficient semantic retrieval; in contrast when the DAN-B network was closer to the heteromodal end of the principal gradient in RH compared with LH, participants showed better visual reasoning on a progressive matrices task. Finally, we established that Control-B dissociates from DAN-B in the effect of verbal task demands on activation in LH and RH. Control-B shows a stronger response to semantic control demands than to working memory load in LH compared with RH, suggesting that lateralised networks near the DMN apex of the principal gradient support controlled semantic retrieval states.

To date, only one previous study has attempted to describe hemispheric differences in the principal gradient (Liang et al., 2021). Despite important differences in methodology, our findings align with Liang et al.’s study: both investigations found higher gradient values in LH than RH for ventromedial prefrontal cortex, IFG and lateral ATL. However, Liang et al. used Yeo’s (2011) 7-network rather than 17-network parcellation, and they extracted separate gradients for LH and RH; consequently, they could not identify the sub-network hemispheric differences that we observed, or directly compare LH and RH networks within the same decomposition. The study by Liang et al. also did not assess the functional consequences of hemispheric differences on the principal gradient, which was the main focus of the current study.

We found that LH parcels were, in general, closer than RH parcels to the DMN apex of the principal gradient, helping to explain why key heteromodal functions – such as language and semantic cognition – are left-lateralised. Margulies et al. (2016) found that the terms language:syntax and language:semantics were among the BrainMap behaviour terms closest to the heteromodal end of the principal gradient; similarly, verbal semantics was towards the heteromodal apex in Neurosynth (a meta-analytic tool; Yarkoni et al., 2011). Language and semantics both depend on the retrieval of heteromodal representations – extracted from diverse sensory-motor features when we acquire concepts and words; moreover, they both require retrieval to be controlled to fit rapidly changing goals and contexts. These different components of semantic cognition – conceptual representations plus control processes – are lateralised to different degrees (Gonzalez Alam et al., 2019). Semantic control processes are supported by a strongly left-lateralised network, encompassing left inferior frontal gyrus, and left posterior middle and inferior temporal cortex (Jackson, 2021; Noonan et al., 2013). The resting-state functional connectivity between these semantic control sites is stronger in the LH compared with the RH (Gonzalez Alam et al., 2019). In contrast, heteromodal conceptual representation is thought to be supported by bilateral ventral ATL (Ding et al., 2020; Lambon Ralph et al., 2017; Patterson et al., 2007). Evidence for bilateral conceptual representation in ventral ATL is provided by neuroimaging studies (Bright et al., 2004; Tranel et al., 2005; Vandenberghe et al., 1996; Visser et al., 2009; Visser and Lambon Ralph, 2011) and neuropsychology; patients with bilateral ventrolateral ATL damage show severe semantic impairment (for example, in semantic dementia), while patients with unilateral lesions have milder deficits (Rice et al., 2018).

This difference between strongly lateralised semantic control processes and bilateral conceptual representations may help to explain why Limbic-A, centred on the ventral anterior temporal lobe, was situated closer to the DMN end of the principal gradient in RH compared with LH. Gonzalez Alam et al. (2019) found that right ATL was more connected to core DMN regions, including angular gyrus and dorsomedial prefrontal cortex; in contrast, left ATL was more connected to left-lateralised sites implicated in semantic control, including left intraparietal sulcus and left anterior insula bordering ventral parts of inferior frontal gyrus. In LH, the principal gradient captures the order of networks from DMN, through the semantic control network, to executive regions (Wang et al., 2020). As a consequence, this proximity (and shared connectivity) of left ATL to semantic control regions might explain the unique gradient difference in Limbic-A. RH components of this network might be closer to the heteromodal apex of the principle gradient because they are further from left-lateralised control networks situated towards the middle of the gradient.

The left-lateralised semantic control network is thought to be partially distinct from multiple demand cortex that responds to executive demands across domains: for example, effects of semantic but not non-semantic control demands are observed in anterior aspects of inferior frontal gyrus and posterior middle temporal gyrus (Davey et al., 2016, 2015; Hoffman et al., 2010; Jackson, 2021; Noonan et al., 2013; Whitney et al., 2012, 2011). Similarly, the frontoparietal control network, defined through analyses of intrinsic functional connectivity, shows a bipartite organisation (Dixon et al., 2018), overlapping with Control-A and Control-B networks within the Yeo et al. (2011) parcellation used in this study. Dixon et al.’s (2018) control subnetwork including more anterior parts of both inferior prefrontal cortex and middle temporal gyrus has a topographical distribution that is similar to the functionally-defined semantic control network (Jackson, 2021; Noonan et al., 2013), and shows stronger interactions with DMN regions than the other control subnetwork. Similarly, the functionally-defined semantic control network shows relatively strong intrinsic connectivity to both DMN, associated with heteromodal integration or abstraction, and domain-general executive and attention networks (Davey et al., 2016). This pattern of connectivity may allow states of controlled semantic cognition in which ongoing activation within DMN regions is shaped through the application of goal representations within executive cortex to promote more weakly-encoded aspects of knowledge (Wang et al., 2020). This finding is consistent with our observation of more efficient semantic cognition when the Control-B network was closer on the principal gradient to DMN in LH as opposed to RH. Gradient differences between the two hemispheres might allow one control subnetwork to connect more strongly with DMN, supporting semantic control in the left hemisphere, while the other control subnetwork in RH connects more strongly with sensory-motor regions, with advantages for demanding tasks that are oriented towards external sensory-motor features. This possibility is consistent with Wang et al. (2014) who found that control network regions in LH have stronger connectivity with DMN, while RH control sites are closer in connectivity to attentional networks.

Like the frontoparietal regions linked to cognitive control, DMN also has subnetworks; this study provides some evidence that these subdivisions within control and DMN networks are functionally related. Just as we found a control network that was closer to the heteromodal end of the principal gradient in LH, DMN-B (the adjacent network), showed the same pattern. DMN-B includes regions such as lateral ATL, angular gyrus, inferior frontal gyrus and dorsomedial prefrontal cortex that are associated with semantic processing in the left hemisphere (Jackson, 2021; Jefferies, 2013; Lambon Ralph et al., 2017; Noonan et al., 2013; Rice et al., 2015), and this DMN variant has repeatedly shown functional dissociations with core DMN regions such as posterior cingulate cortex and more ventromedial prefrontal regions (Chiou et al., 2020; Zhang et al., 2020; referred to here as DMN-A). DMN-B is associated with lateralised cognitive processes, like language and semantics, as well as social cognition (Andrews-Hanna et al., 2014). This network shows responses to externally-generated, conceptual tasks, including those that interface with perception. In contrast, DMN-A or core DMN is thought to be more detached from perception, and is engaged by internally generated, self-referential and autobiographical memory processing (Chiou et al., 2020). It is interesting to note that it is DMN-B, not core DMN, that shows a lateralised position on the principal gradient. This is consistent with the possibility that lateralisation reflects the need to sustain and/or control heteromodal semantic retrieval (as opposed to the need to support internally-generated mental states, which are also associated with the heteromodal end of the principal gradient).

We also found evidence of significant differences in lateralisation patterns within attentional networks, with both DAN and VAN falling closer to the heteromodal end of the gradient in LH. Although attention has been traditionally conceptualised as a right-lateralised cognitive function, contemporary neuroscientific research paints a more nuanced picture with complex patterns of lateralisation across the traditionally accepted ventral and dorsal attention networks (Corbetta and Shulman, 2002; Jeong and Xu, 2016; Szczepanski et al., 2010; Thiebaut de Schotten et al., 2011a, 2011b). Critically, the DAN also plays a role in the flexible coupling of the control network across hemispheres and subdivisions (Dixon et al., 2018; Wang et al., 2014). Both DAN and control networks showed significant but opposing behavioural associations in our individual differences analysis of the position of networks on the principal gradient across hemispheres. Hemispheric differences in DAN-B were related to Raven’s matrices performance, but in contrast to semantic cognition, participants whose DAN-B was closer to the heteromodal end of the gradient in RH were better at the task. RH is particularly activated during the performance of this task (Bishop et al., 2008; Haier et al., 1988; Prabhakaran et al., 1997). Moreover, previous research has linked performance on progressive matrices to attentional capacity (Schweizer and Moosbrugger, 2004), and performance can also be decomposed into in two components relating to perceptual and executive attention (Ren et al., 2012), with the latter corresponding more closely to the DAN (Corbetta et al., 2008; Corbetta and Shulman, 2002) and accounting for more variance in visual reasoning tasks like progressive matrices (Ren et al., 2013).

There are several limitations of the current study. Our methods did not allow us to investigate the source of the network asymmetries at the sub-network or parcel level, since the choice of parcellation (Schaefer et al., 2018) does not provide homotopic regions that can be compared. Future research could address this by using parcellations that are suitable for determining homotopy (Glasser et al., 2016). Also, it remains unclear why attentional networks (DAN-A; DAN-B and VAN-A) were closer to the heteromodal end of the principal gradient in LH, even when the opposite pattern for DAN-B (closer proximity to heteromodal cortex in RH) was associated with better visual attention. One possibility is that these attention networks can also support controlled semantic cognition, to varying degrees across people, and that these patterns of left-lateralised and right-lateralised connectivity are in competition. Future research could test whether the position of networks along the principal gradient relates to their capacity for efficient interaction, and whether there are differences in physical distance along the cortical surface in the two hemispheres that reflect the connectivity gradient differences we described. Future studies could also investigate control processes in more cognitive domains to determine the functional basis of the semantic-to-visual reasoning dissociation that we observed.

## 5. Conclusions

We found that networks associated with higher-order cognition in LH are positioned closer to the heteromodal end of the principal gradient, including the DMN, control, limbic and attentional networks; in contrast, there were no differences in sensorimotor networks, in line with the literature on functional homotopy. The control-DAN dissociation we observed is compatible with recent proposals of a “inward-outward” organisational principle for control networks that differs across the hemispheres, with a privileged interaction of DMN-B and control-B in LH (Dixon et al., 2018; Wang et al., 2014). Individual difference analysis showed that relative network position across the hemispheres has functional consequences in the efficient implementation of lateralised cognitive processes: proximity of DMN to control regions in LH predicted more efficient semantic processing, while proximity of DAN to control regions in RH predicted better visual reasoning.

## Declarations

### Availability of data and material

Gradient maps one to ten from the group-averaged dimension reduction analysis described in section 2.3.2. below are publicly available on NeuroVault in a collection (https://neurovault.org/collections/6746/). Raw fMRI and questionnaire data are restricted in accordance with ERC and EU regulations.

### Code availability

All code used in the production of this manuscript is available from the corresponding authors upon reasonable request.

### Ethics approval

The study was approved by the York Neuroimaging Centre Ethics Committee. This study was performed in line with the principles of the Declaration of Helsinki.

### Consent to participate

Written informed consent was obtained for all participants.

### Consent for publication

Consent for publication of anonymised, group-level data was provided as part of the written informed consent.

## Acknowledgments

None.

## SUPPLEMENTARY MATERIALS

**Supplementary Figure 1.**
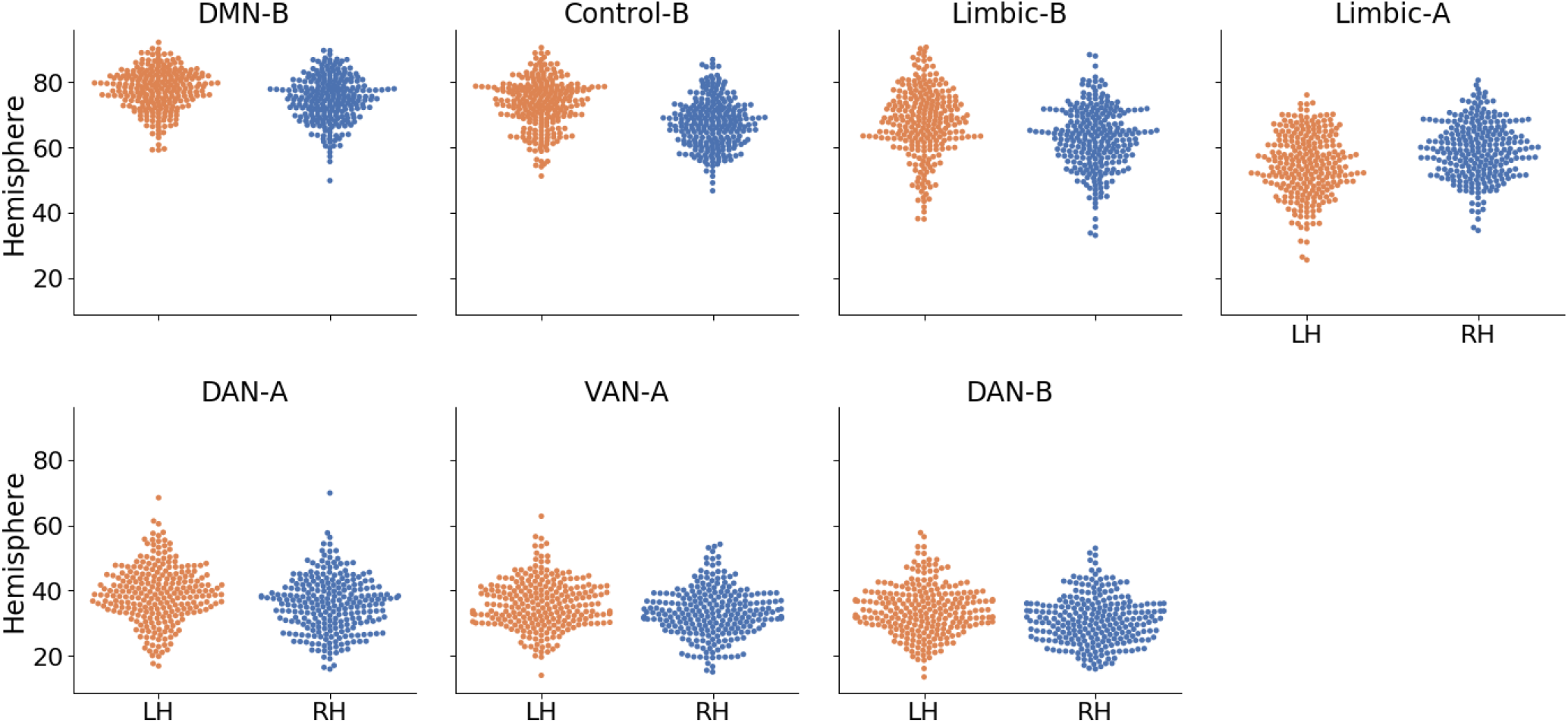
Breakdown of networks that show hemispheric differences for Gradient 1.Supplementary Analysis: Gradient 2

**Supplementary Figure 2.**
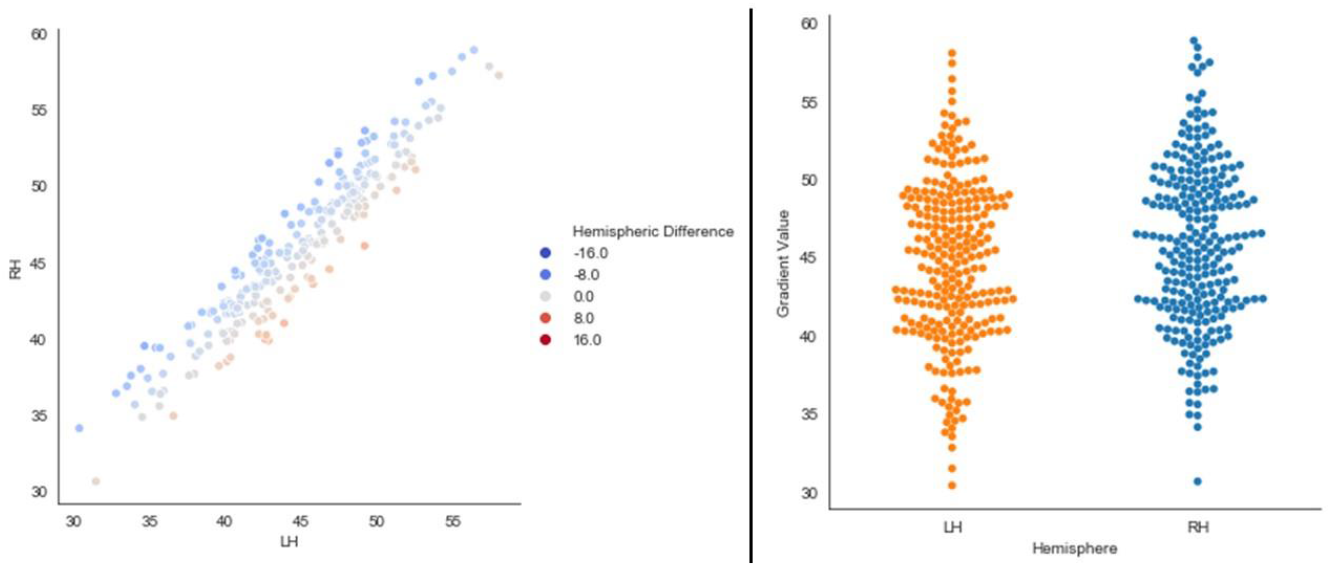
The left panel depicts a linear relationship in our sample’s mean left and right hemisphere values in the second gradient. The right panel depicts the distributions of these mean global hemispheric values per participant in our sample. The ‘Hemispheric Difference’ legend of the scatterplot depicts the result of subtracting the L – R gradient loadings for the whole hemisphere per participant (negative values show right bias).

**Supplementary Figure 3.**
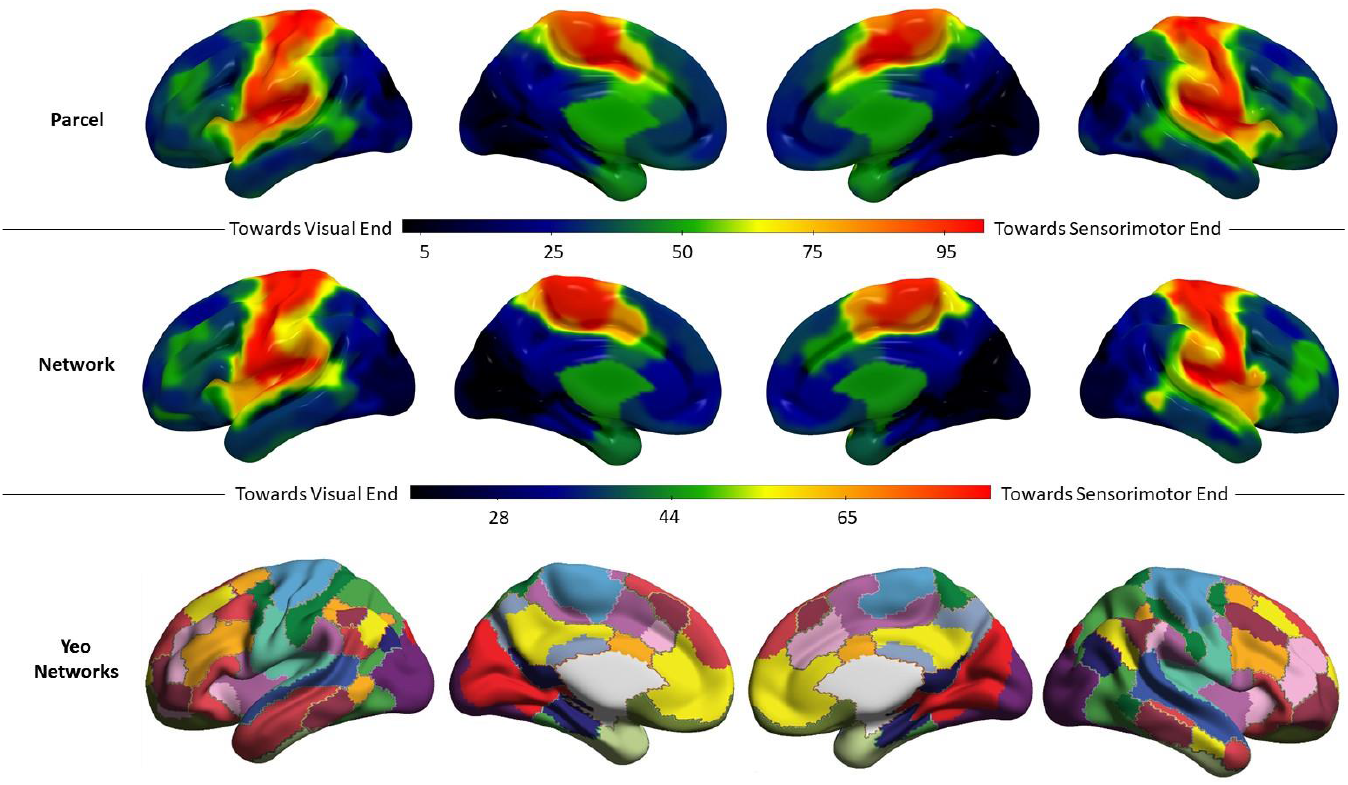
Top row: Group mean second gradient value for each parcel in Schaeffer’s 400-parcel solution for our sample of 254 participants. Middle row: Group mean second gradient value for each network in Yeo’s 17-network solution for our sample of 254 participants. Gradient units are arbitrary and have been normalised on a 0-100 minmax scale. Bottom row: 17 network parcellation by Yeo et al. The colour code followed in this figure replicates that of Buckner et al. (2011)

**Supplementary Figure 4.**
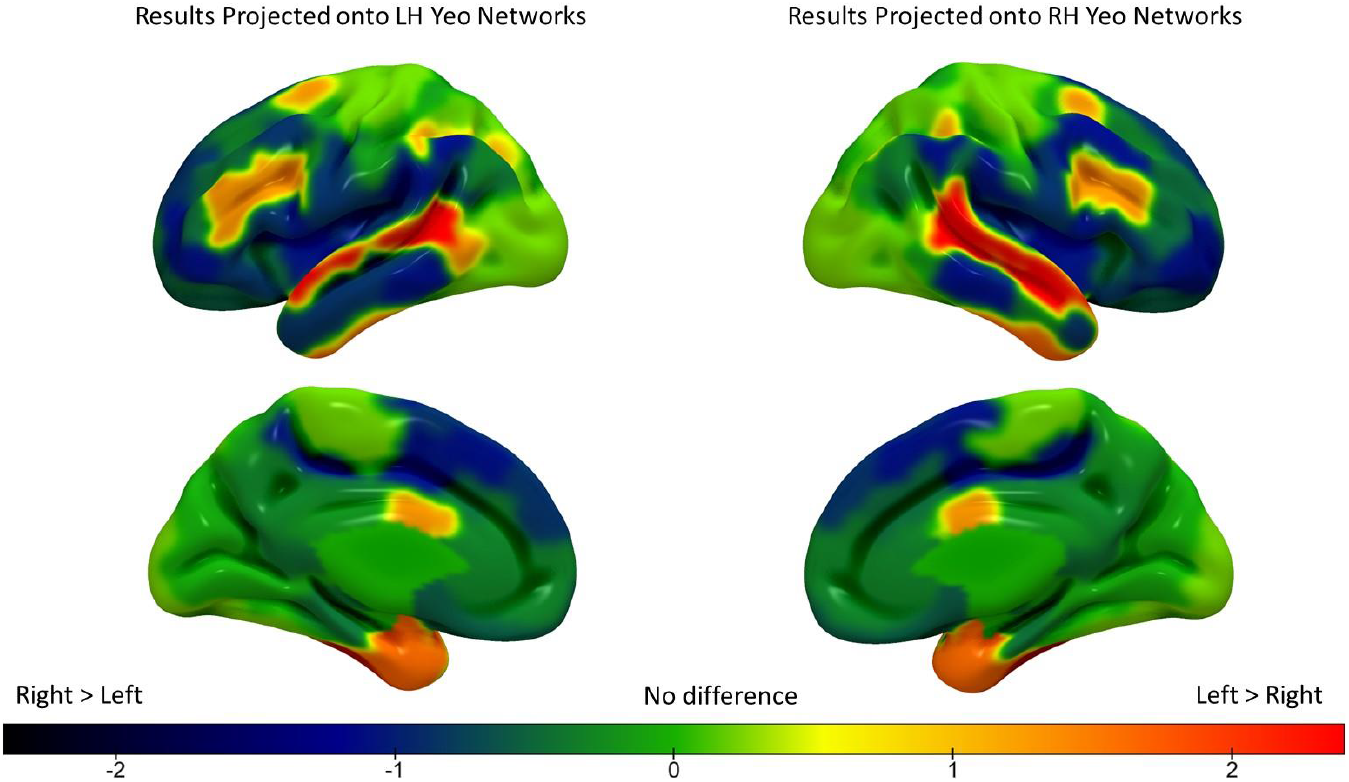
Hemispheric differences in second gradient values across the 17 Yeo networks (z-scored). The warm colours represent a greater principal gradient loading in the LH compared to RH, and the cool colours represent a greater principal gradient loading in RH compared to LH.

**Supplementary Figure 5.**
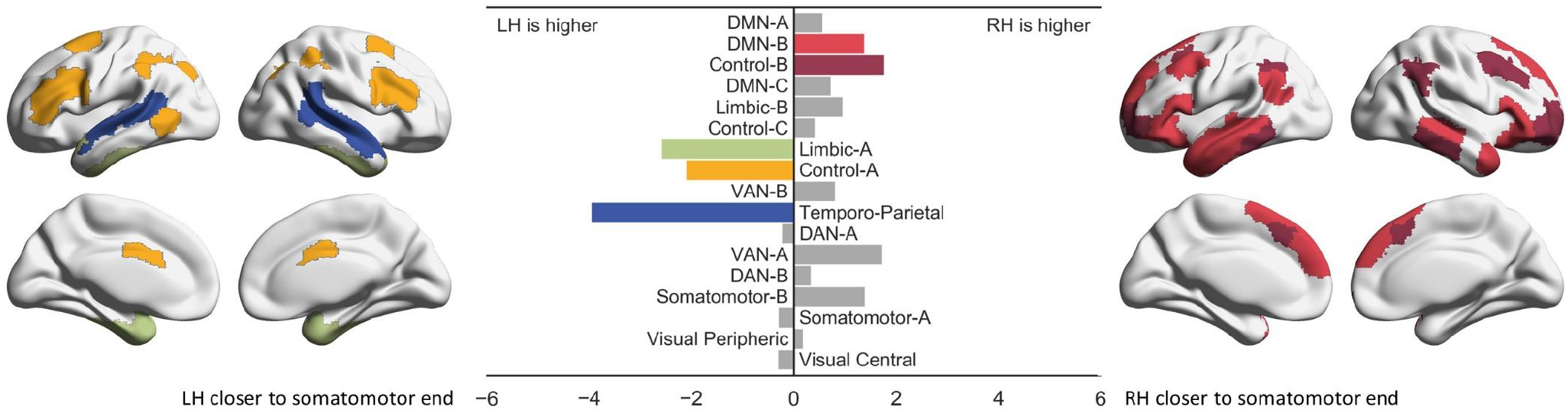
Results of pairwise bootstrapped replicates permutation testing of LH vs RH means for each network (5,000 simulations, Bonferroni-corrected alpha for 17 comparisons equal to 0.00294). The size of each bar reflects the normalized (0-100) empirically observed mean difference across the hemispheres for each network. Coloured bars denote networks that showed significant differences and are colour-coded to the significant networks in the brain maps.

**Table S1.** OLS Regression results for semantic task contrasts.

| **Network** | **Contrast** | **$\beta$** | **P (uncorrected)** |
| --- | --- | --- | --- |
| Control-B | Word > Picture | .09 | .23 |
|  | Strong > Weak | -.06 | .55 |
| Default-B | Word > Picture | .08 | .26 |
|  | Strong > Weak | .01 | .87 |
| DAN-A | Word > Picture | .04 | .62 |
|  | Strong > Weak | .11 | .21 |
| DAN-B | Word > Picture | .05 | .52 |
|  | Strong > Weak | .11 | .2 |
| Limbic-B | Word > Picture | -.01 | .88 |
|  | Strong > Weak | -.08 | .42 |
| Limbic-A | Word > Picture | .05 | .55 |
|  | Strong > Weak | .15 | .11 |
| VAN-A | Word > Picture | .01 | .89 |
|  | Strong > Weak | .01 | .89 |

## References

Andrews-Hanna, J.R., Smallwood, J., Spreng, R.N., 2014. The default network and self-generated thought: component processes, dynamic control, and clinical relevance. Annals of the New York Academy of Sciences 1316, 29–52. https://doi.org/https://doi.org/10.1111/nyas.12360

Badre, D., 2008. Cognitive control, hierarchy, and the rostro-caudal organization of the frontal lobes.Trends in Cognitive Sciences 12, 193–200. https://doi.org/10.1016/j.tics.2008.02.004

Badre, D., D’Esposito, M., 2007. Functional Magnetic Resonance Imaging Evidence for a Hierarchical Organization of the Prefrontal Cortex. Journal of Cognitive Neuroscience 19, 2082–2099. https://doi.org/10.1162/jocn.2007.19.12.2082

Bajada, C.J., Jackson, R.L., Haroon, H.A., Azadbakht, H., Parker, G.J.M., Lambon Ralph, M.A., Cloutman, L.L., 2017. A graded tractographic parcellation of the temporal lobe. NeuroImage 155, 503–512. https://doi.org/10.1016/j.neuroimage.2017.04.016

Bajada, C.J., Trujillo-Barreto, N.J., Parker, G.J.M.M., Cloutman, L.L., Lambon Ralph, M.A., 2019. A structural connectivity convergence zone in the ventral and anterior temporal lobes: Data-driven evidence from structural imaging. Cortex 120, 298–307. https://doi.org/ https://doi.org/10.1016/j.cortex.2019.06.014

Bartolomeo, P., Seidel Malkinson, T., 2019. Hemispheric lateralization of attention processes in the human brain. Current Opinion in Psychology 29, 90–96. https://doi.org/10.1016/j.copsyc.2018.12.023

Behzadi, Y., Restom, K., Liau, J., Liu, T.T., 2007. A component based noise correction method (CompCor) for BOLD and perfusion based fMRI. NeuroImage 37, 90–101. https://doi.org/10.1016/j.neuroimage.2007.04.042

Bishop, S.J., Fossella, J., Croucher, C.J., Duncan, J., 2008. COMT val158met genotype affects recruitment of neural mechanisms supporting fluid intelligence. Cerebral Cortex 18, 2132–2140. https://doi.org/10.1093/cercor/bhm240

Bressler, S.L., Menon, V., 2010. Large-scale brain networks in cognition: emerging methods and principles. Trends in Cognitive Sciences 14, 277–290. https://doi.org/10.1016/j.tics.2010.04.004

Bright, P., Moss, H., Tyler, L.K., 2004. Unitary vs multiple semantics: PET studies of word and picture processing. Brain and Language 89, 417–432. https://doi.org/10.1016/j.bandl.2004.01.010

Buckner, R.L., Krienen, F.M., Castellanos, A., Diaz, J.C., Yeo, B.T.T., 2011. The organization of the human cerebellum estimated by intrinsic functional connectivity. Journal of neurophysiology 106, 2322–2345. https://doi.org/10.1152/jn.00339.2011

Chiou, R., Humphreys, G.F., Lambon Ralph, M.A., 2020. Bipartite functional fractionation within the default network supports disparate forms of internally oriented cognition. Cerebral Cortex 30, 5484–5501. https://doi.org/10.1093/cercor/bhaa130

Coltheart, M., 1981. The MRC psycholinguistic database. The Quarterly Journal of Experimental Psychology Section A 33, 497–505. https://doi.org/10.1080/14640748108400805

Corbetta, M., Patel, G., Shulman, G.L., 2008. The Reorienting System of the Human Brain: From Environment to Theory of Mind. Neuron 58, 306–324. https://doi.org/10.1016/j.neuron.2008.04.017

Corbetta, M., Shulman, G.L., 2002. Control of Goal-Directed and Stimulus-Driven Attention in the Brain. Nature Reviews Neuroscience 3, 215–229. https://doi.org/10.1038/nrn755

Davey, J., Cornelissen, P.L., Thompson, H.E., Sonkusare, S., Hallam, G., Smallwood, J., Jefferies, E., 2015. Automatic and Controlled Semantic Retrieval: TMS Reveals Distinct Contributions of Posterior Middle Temporal Gyrus and Angular Gyrus. The Journal of neuroscience : the official journal of the Society for Neuroscience 35, 15230–9. https://doi.org/10.1523/JNEUROSCI.4705-14.2015

Davey, J., Thompson, H.E., Hallam, G., Karapanagiotidis, T., Murphy, C., de Caso, I., Krieger-Redwood, K., Bernhardt, B.C., Smallwood, J., Jefferies, E., 2016. Exploring the role of the posterior middle temporal gyrus in semantic cognition: Integration of anterior temporal lobe with executive processes. NeuroImage 137, 165–177. https://doi.org/10.1016/j.neuroimage.2016.05.051

Ding, J., Chen, K., Liu, H., Huang, L., Chen, Y., Lv, Y., Yang, Q., Guo, Q., Han, Z., Lambon Ralph, M.A., 2020. A unified neurocognitive model of semantics language social behaviour and face recognition in semantic dementia. Nature Communications 11, 1–14. https://doi.org/10.1038/s41467-020-16089-9

Dixon, M.L., de La Vega, A., Mills, C., Andrews-Hanna, J., Spreng, R.N., Cole, M.W., Christoff, K., 2018. Heterogeneity within the frontoparietal control network and its relationship to the default and dorsal attention networks. Proceedings of the National Academy of Sciences 115, E1598–E1607. https://doi.org/10.1073/pnas.1715766115

Evans, M., Krieger-Redwood, K., Gonzalez Alam, T.R.J., Smallwood, J., Jefferies, E., 2020. Controlled semantic summation correlates with intrinsic connectivity between default mode and control networks. Cortex 129, 356–375. https://doi.org/ https://doi.org/10.1016/j.cortex.2020.04.032

Fuster, J.M., 2001. The prefrontal cortex - An update: Time is of the essence. Neuron 30, 319–333. https://doi.org/10.1016/S0896-6273(01)00285-9

Gao, Z., Zheng, L., Chiou, R., Gouws, A., Krieger-Redwood, K., Wang, X., Varga, D., Lambon Ralph, M.A., Smallwood, J., Jefferies, E., 2020. Distinct and Common Neural Coding of Semantic and Non-semantic Control Demands. bioRxiv 2020.11.16.384883. https://doi.org/10.1101/2020.11.16.384883

Glasser, M.F., Coalson, T.S., Robinson, E.C., Hacker, C.D., Harwell, J., Yacoub, E., Ugurbil, K., Andersson, J., Beckmann, C.F., Jenkinson, M., Smith, S.M., van Essen, D.C., 2016. A multi-modal parcellation of human cerebral cortex. Nature 536, 171–178. https://doi.org/10.1038/nature18933

Gonzalez Alam, T., Murphy, C., Smallwood, J., Jefferies, E., 2018. Meaningful inhibition: Exploring the role of meaning and modality in response inhibition. NeuroImage 181, 108–119. https://doi.org/10.1016/j.neuroimage.2018.06.074

Gonzalez Alam, T.R. del J., Karapanagiotidis, T., Smallwood, J., Jefferies, E., 2019. Degrees of lateralisation in semantic cognition: Evidence from intrinsic connectivity. NeuroImage 202, 116089. https://doi.org/10.1016/j.neuroimage.2019.116089

Gonzalez Alam, T.RJ., Krieger-Redwood, K., Evans, M., Rice, G.E., Smallwood, J., Jefferies, E., 2021. Intrinsic connectivity of anterior temporal lobe relates to individual differences in semantic retrieval for landmarks. Cortex 134, 76–91. https://doi.org/10.1016/j.cortex.2020.10.007

Gotts, S.J., Jo, H.J., Wallace, G.L., Saad, Z.S., Cox, R.W., Martin, A., 2013. Two distinct forms of functional lateralization in the human brain. Proceedings of the National Academy of Sciences of the United States of America 110, E3435–44. https://doi.org/10.1073/pnas.1302581110

Haier, R.J., Siegel, B. v, Nuechterlein, K.H., Hazlett, E., Wu, J.C., Paek, J., Browning, H.L., Buchsbaum, M.S., 1988. Cortical glucose metabolic rate correlates of abstract reasoning and attention studied with positron emission tomography. Intelligence 12, 199–217. https://doi.org/https://doi.org/10.1016/0160-2896(88)90016-5

Hearne, L.J., Cocchi, L., Zalesky, A., Mattingley, J.B., 2017. Reconfiguration of brain network architectures between resting-state and complexity-dependent cognitive reasoning. Journal of Neuroscience 37, 8399–8411. https://doi.org/10.1523/JNEUROSCI.0485-17.2017

Hoffman, P., Jefferies, E., Lambon Ralph, M. a, 2010. Ventrolateral prefrontal cortex plays an executive regulation role in comprehension of abstract words: convergent neuropsychological and repetitive TMS evidence. The Journal of Neuroscience 30, 15450–15456. https://doi.org/10.1523/JNEUROSCI.3783-10.2010

Huntenburg, J.M., Bazin, P.L., Margulies, D.S., 2018. Large-Scale Gradients in Human Cortical Organization. Trends in Cognitive Sciences 22, 21–31. https://doi.org/10.1016/j.tics.2017.11.002

Jackson, R.L., 2021. The neural correlates of semantic control revisited. NeuroImage 224, 117444. https://doi.org/10.1016/j.neuroimage.2020.117444

Jackson, R.L., Bajada, C.J., Ralph, M.A.L., Cloutman, L.L., 2019. The Graded Change in Connectivity across the Ventromedial Prefrontal Cortex Reveals Distinct Subregions. Cerebral Cortex 1–16. https://doi.org/10.1093/cercor/bhz079

Jackson, R.L., Bajada, C.J., Rice, G.E., Cloutman, L.L., Lambon Ralph, M.A., 2017. An emergent functional parcellation of the temporal cortex. NeuroImage 1–15. https://doi.org/10.1016/j.neuroimage.2017.04.024

Jefferies, E., 2013. The neural basis of semantic cognition: Converging evidence from neuropsychology, neuroimaging and TMS. Cortex 49, 611–625. https://doi.org/10.1016/j.cortex.2012.10.008

Jeong, S.K., Xu, Y., 2016. The impact of top-down spatial attention on laterality and hemispheric asymmetry in the human parietal cortex. Journal of Vision 16, 1–21. https://doi.org/10.1167/16.10.2

Jo, H.J., Saad, Z.S., Gotts, S.J., Martin, A., Cox, R.W., 2012. Quantifying Agreement between Anatomical and Functional Interhemispheric Correspondences in the Resting Brain. PLoS ONE 7. https://doi.org/10.1371/journal.pone.0048847

Joliot, M., Tzourio-Mazoyer, N., Mazoyer, B., 2016. Intra-hemispheric intrinsic connectivity asymmetry and its relationships with handedness and language Lateralization. Neuropsychologia 93, 437–447. https://doi.org/10.1016/j.neuropsychologia.2016.03.013

Karapanagiotidis, T., Bernhardt, B.C., Jefferies, E., Smallwood, J., 2017. Tracking thoughts: Exploring the neural architecture of mental time travel during mind-wandering. NeuroImage 147, 272–281. https://doi.org/10.1016/j.neuroimage.2016.12.031

Karapanagiotidis, T., Vidaurre, D., Quinn, A.J., Vatansever, D., Poerio, G.L., Turnbull, A., Ho, N.S.P., Leech, R., Bernhardt, B.C., Jefferies, E., Margulies, D.S., Nichols, T.E., Woolrich, M.W., Smallwood, J., 2020. The psychological correlates of distinct neural states occurring during wakeful rest. Scientific Reports 10, 21121. https://doi.org/10.1038/s41598-020-77336-z

Karolis, V.R., Corbetta, M., Thiebaut de Schotten, M., 2019. The architecture of functional lateralisation and its relationship to callosal connectivity in the human brain. Nature Communications 10, 1417. https://doi.org/10.1038/s41467-019-09344-1

Knecht, S., Deppe, M., Dräger, B., Bobe, L., Lohmann, H., Ringelstein, E.B., Henningsen, H., 2000. Language lateralization in healthy right-handers. Brain 123, 74–81. https://doi.org/10.1093/brain/123.1.74

Koechlin, E., Ody, C., Kouneiher, F., 2003. The Architecture of Cognitive Control in the Human Prefrontal Cortex. Science 302, 1181–1185. https://doi.org/10.1126/science.1088545

Krieger-Redwood, K., Teige, C., Davey, J., Hymers, M., Jefferies, E., 2015. Conceptual control across modalities: graded specialisation for pictures and words in inferior frontal and posterior temporal cortex. Neuropsychologia 76, 92–107. https://doi.org/10.1016/j.neuropsychologia.2015.02.030

Lambon Ralph, M.A., Jefferies, E., Patterson, K., Rogers, T.T., 2017. The neural and computational bases of semantic cognition. Nature Reviews Neuroscience 18, 42–55. https://doi.org/10.1038/nrn.2016.150

Liang, X., Zhao, C., Jin, X., Jiang, Y., Yang, L., Chen, Y., Gong, G., 2021. Sex-related human brain asymmetry in hemispheric functional gradients. NeuroImage 229, 117761. https://doi.org/ https://doi.org/10.1016/j.neuroimage.2021.117761

Mancuso, L., Costa, T., Nani, A., Manuello, J., Liloia, D., Gelmini, G., Panero, M., Duca, S., Cauda, F., 2019. The homotopic connectivity of the functional brain: a meta-analytic approach. Scientific Reports 9, 1–19. https://doi.org/10.1038/s41598-019-40188-3

Margulies, D.S., Ghosh, S.S., Goulas, A., Falkiewicz, M., Huntenburg, J.M., Langs, G., Bezgin, G., Eickhoff, S.B., Castellanos, F.X., Petrides, M., Jefferies, E., Smallwood, J., 2016. Situating the default-mode network along a principal gradient of macroscale cortical organization. Proceedings of the National Academy of Sciences of the United States of America 113, 12574–12579. https://doi.org/10.1073/pnas.1608282113

Mckeown, B., Strawson, W.H., Wang, H.T., Karapanagiotidis, T., Vos de Wael, R., Benkarim, O., Turnbull, A., Margulies, D., Jefferies, E., McCall, C., Bernhardt, B., Smallwood, J., 2020. The relationship between individual variation in macroscale functional gradients and distinct aspects of ongoing thought. NeuroImage 220. https://doi.org/10.1016/j.neuroimage.2020.117072

Medaglia, J.D., Lynall, M.-E., Bassett, D.S., 2015. Cognitive Network Neuroscience. Journal of Cognitive Neuroscience 27, 1471–1491. https://doi.org/10.1162/jocn_a_00810

Mikolov, T., Sutskever, I., Chen, K., Corrado, G.S., Dean, J., 2013. Distributed Representations of Words and Phrases and their Compositionality, in: Burges, C.J.C., Bottou, L., Welling, M., Ghahramani, Z., Weinberger, K.Q. (Eds.), Advances in Neural Information Processing Systems. Curran Associates, Inc., pp. 3111–3119.

Niendam, T.A., Laird, A.R., Ray, K.L., Dean, Y.M., Glahn, D.C., Carter, C.S., 2012. Meta-analytic evidence for a superordinate cognitive control network subserving diverse executive functions. Cognitive, Affective, & Behavioral Neuroscience 12, 241–268. https://doi.org/10.3758/s13415-011-0083-5

Noonan, K.A., Jefferies, E., Visser, M., Lambon Ralph, M.A., 2013. Going beyond Inferior Prefrontal Involvement in Semantic Control: Evidence for the Additional Contribution of Dorsal Angular Gyrus and Posterior Middle Temporal Cortex. Journal of Cognitive Neuroscience 25, 1824–50. https://doi.org/10.1162/jocn

Paquola, C., Vos De Wael, R., Wagstyl, K., Bethlehem, R.A.I., Hong, S.J., Seidlitz, J., Bullmore, E.T., Evans, A.C., Misic, B., Margulies, D.S., Smallwood, J., Bernhardt, B.C., 2018. Microstructural and Functional Gradients are Increasingly Dissociated in Transmodal Cortices. bioRxiv 1–28. https://doi.org/10.1101/488700

Patterson, K., Nestor, P.J., Rogers, T.T., 2007. Where do you know what you know? The representation of semantic knowledge in the human brain. Nature Reviews Neuroscience 8, 976–987. https://doi.org/10.1038/nrn2277

Petrides, M., 2005. Lateral prefrontal cortex: architectonic and functional organization. Philosophical Transactions of the Royal Society B: Biological Sciences 360, 781–795. https://doi.org/10.1098/rstb.2005.1631

Poerio, G.L., Sormaz, M., Wang, H.T., Margulies, D., Jefferies, E., Smallwood, J., 2017. The role of the default mode network in component processes underlying the wandering mind. Social Cognitive and Affective Neuroscience 12, 1047–1062. https://doi.org/10.1093/scan/nsx041

Power, J.D., Mitra, A., Laumann, T.O., Snyder, A.Z., Schlaggar, B.L., Petersen, S.E., 2014. Methods to detect, characterize, and remove motion artifact in resting state fMRI. NeuroImage 84, 320–341. https://doi.org/10.1016/j.neuroimage.2013.08.048

Prabhakaran, V., Smith, J.A.L., Desmond, J.E., Glover, G.H., Gabrieli, J.D.E., 1997. Neural substrates of fluid reasoning: An fMRI study of neocortical activation during performance of the Raven’s Progressive Matrices Test. Cognitive Psychology 33, 43–63. https://doi.org/10.1006/cogp.1997.0659

Raemaekers, M., Schellekens, W., Petridou, N., Ramsey, N.F., 2018. Knowing left from right: asymmetric functional connectivity during resting state. Brain Structure and Function 223, 1909–1922. https://doi.org/10.1007/s00429-017-1604-y

Raven, J., Matrices, R.P., Hill, M., Scales, V., 1994. Manual for Raven’s progressive matrices and mill hill vocabulary scales. Advanced progressive matrices.

Ren, X., Goldhammer, F., Moosbrugger, H., Schweizer, K., 2012. How does attention relate to the ability-specific and position-specific components of reasoning measured by APM? Learning and Individual Differences 22, 1–7. https://doi.org/10.1016/j.lindif.2011.09.009

Ren, X., Schweizer, K., Xu, F., 2013. The sources of the relationship between sustained attention and reasoning. Intelligence 41, 51–58. https://doi.org/10.1016/j.intell.2012.10.006

Rice, G.E., Caswell, H., Moore, P., Ralph, M.A.L., Hoffman, P., 2018. Revealing the Dynamic Modulations That Underpin a Resilient Neural Network for Semantic Cognition: An fMRI Investigation in Patients With Anterior Temporal Lobe Resection. Cerebral Cortex 28, 3004–3016. https://doi.org/10.1093/cercor/bhy116

Rice, G.E., Lambon Ralph, M.A., Hoffman, Paul., 2015. The Roles of Left Versus Right Anterior Temporal Lobes in Conceptual Knowledge: An ALE Meta-analysis of 97 Functional Neuroimaging Studies. Cerebral Cortex 25, 4374–4391. https://doi.org/10.1093/cercor/bhv024

Schaefer, A., Kong, R., Gordon, E.M., Laumann, T.O., Zuo, X.-N., Holmes, A.J., Eickhoff, S.B., Yeo, B.T.T., 2018. Local-Global Parcellation of the Human Cerebral Cortex from Intrinsic Functional Connectivity MRI. Cerebral Cortex 28, 3095–3114. https://doi.org/10.1093/cercor/bhx179

Schweizer, K., Moosbrugger, H., 2004. Attention and working memory as predictors of intelligence. Intelligence 32, 329–347. https://doi.org/10.1016/j.intell.2004.06.006

Sormaz, M., Murphy, C., Wang, H., Hymers, M., Poerio, G., Margulies, D.S., Sormaz, M., Murphy, C., Wang, H., Hymers, M., Karapanagiotidis, T., Poerio, G., 2018. Default mode network can support the level of detail in experience during active task states. Proceedings of the National Academy of Sciences 115, 9318–9323. https://doi.org/10.1073/pnas.1817966115

Spreng, R.N., Sepulcre, J., Turner, G.R., Stevens, W.D., Schacter, D.L., 2013. Intrinsic architecture underlying the relations among the default, dorsal attention, and frontoparietal control networks of the human brain. Journal of cognitive neuroscience 25, 74–86. https://doi.org/10.1162/jocn_a_00281

Stark, D.E., Margulies, D.S., Shehzad, Z.E., Reiss, P., Kelly, A.M.C., Uddin, L.Q., Gee, D.G., Roy, A.K., Banich, M.T., Castellanos, F.X., Milham, M.P., 2008. Regional variation in interhemispheric coordination of intrinsic hemodynamic fluctuations. The Journal of neuroscience : the official journal of the Society for Neuroscience 28, 13754–13764. https://doi.org/10.1523/JNEUROSCI.4544-08.2008

Szczepanski, S.M., Konen, C.S., Kastner, S., 2010. Mechanisms of spatial attention control in frontal and parietal cortex. Journal of Neuroscience 30, 148–160. https://doi.org/10.1523/JNEUROSCI.3862-09.2010

Thiebaut de Schotten, M., Dell’Acqua, F., Forkel, S.J., Simmons, A., Vergani, F., Murphy, D.G.M., Catani, M., 2011a. A lateralized brain network for visuospatial attention. Nature Neuroscience 14, 1245–1246. https://doi.org/10.1038/nn.2905

Thiebaut de Schotten, M., ffytche, D.H., Bizzi, A., Dell’Acqua, F., Allin, M., Walshe, M., Murray, R., Williams, S.C., Murphy, D.G.M., Catani, M., 2011b. Atlasing location, asymmetry and inter-subject variability of white matter tracts in the human brain with MR diffusion tractography. NeuroImage 54, 49–59. https://doi.org/10.1016/j.neuroimage.2010.07.055

Thiebaut De Schotten, M., Urbanski, M., Batrancourt, B., Levy, R., Dubois, B., Cerliani, L., Volle, E., 2017. Rostro-caudal architecture of the frontal lobes in humans. Cerebral Cortex 27, 4033–4047. https://doi.org/10.1093/cercor/bhw215

Tranel, D., Grabowski, T.J., Lyon, J., Damasio, H., 2005. Naming the Same Entities from Visual or from Auditory Stimulation Engages Similar Regions of Left Inferotemporal Cortices. Journal of Cognitive Neuroscience. https://doi.org/10.1162/0898929055002508

Turnbull, A., Wang, H.-T., Schooler, J.W., Jefferies, E., Margulies, D.S., Smallwood, J., 2018. The ebb and flow of attention: Between-subject variation in intrinsic connectivity and cognition associated with the dynamics of ongoing experience. NeuroImage 185, 286–299. https://doi.org/10.1016/j.neuroimage.2018.09.069

van Heuven, W.J.B., Mandera, P., Keuleers, E., Brysbaert, M., 2014. SUBTLEX-UK: A new and improved word frequency database for British English. Quarterly Journal of Experimental Psychology 67, 1176–1190. https://doi.org/10.1080/17470218.2013.850521

Vandenberghe, R., Price, C., Wise, R., Josephs, O., Frackowiak, R.S.J., 1996. Functional anatomy of a common semantic system for words and pictures. Nature 383, 254–256. https://doi.org/10.1038/383254a0

Vatansever, D., Bzdok, D., Wang, H., Mollo, G., Sormaz, M., Murphy, C., Karapanagiotidis, T., Smallwood, J., Jefferies, E., 2017. Varieties of semantic cognition revealed through simultaneous decomposition of intrinsic brain connectivity and behaviour. NeuroImage 158, 1–11. https://doi.org/10.1016/j.neuroimage.2017.06.067

Vincent, J.L., Kahn, I., Snyder, A.Z., Raichle, M.E., Buckner, R.L., 2008. Evidence for a Frontoparietal Control System Revealed by Intrinsic Functional Connectivity. Journal of Neurophysiology 100, 3328–3342. https://doi.org/10.1152/jn.90355.2008

Visser, M., Jefferies, E., Lambon Ralph, M.A., 2009. Semantic Processing in the Anterior Temporal Lobes: A Meta-analysis of the Functional Neuroimaging Literature. Journal of Cognitive Neuroscience 22, 1083–1094. https://doi.org/10.1162/jocn.2009.21309

Visser, M., Lambon Ralph, M. a, 2011. Differential contributions of bilateral ventral anterior temporal lobe and left anterior superior temporal gyrus to semantic processes. Journal of cognitive neuroscience 23, 3121–3131. https://doi.org/10.1162/jocn_a_00007

Vos de Wael, R., Benkarim, O., Paquola, C., Lariviere, S., Royer, J., Tavakol, S., Xu, T., Hong, S.J., Langs, G., Valk, S., Misic, B., Milham, M., Margulies, D., Smallwood, J., Bernhardt, B.C., 2020. BrainSpace: a toolbox for the analysis of macroscale gradients in neuroimaging and connectomics datasets. Communications Biology 3. https://doi.org/10.1038/s42003-020-0794-7

Vos De Wael, R., Larivière, S., Caldairou, B., Hong, S.J., Margulies, D.S., Jefferies, E., Bernasconi, A., Smallwood, J., Bernasconi, N., Bernhardt, B.C., 2018. Anatomical and microstructural determinants of hippocampal subfield functional connectome embedding. Proceedings of the National Academy of Sciences of the United States of America 115, 10154–10159. https://doi.org/10.1073/pnas.1803667115

Wang, D., Buckner, R.L., Liu, H., 2014. Functional Specialization in the Human Brain Estimated By Intrinsic Hemispheric Interaction. Journal of Neuroscience 34, 12341–12352. https://doi.org/10.1523/JNEUROSCi.0787-14.2014

Wang, H.T., Poerio, G., Murphy, C., Bzdok, D., Jefferies, E., Smallwood, J., 2018. Dimensions of Experience: Exploring the Heterogeneity of the Wandering Mind. Psychological Science 29, 56–71. https://doi.org/10.1177/0956797617728727

Wang, X., Bernhardt, B.C., Karapanagiotidis, T., de Caso, I., Gonzalez Alam, T.R. del J., Cotter, Z., Smallwood, J., Jefferies, E., 2018. The structural basis of semantic control: Evidence from individual differences in cortical thickness. NeuroImage 181, 480–489. https://doi.org/10.1016/j.neuroimage.2018.07.044

Wang, X., Margulies, D.S., Smallwood, J., Jefferies, E., 2020. A gradient from long-term memory to novel cognition: Transitions through default mode and executive cortex. NeuroImage 220, 117074. https://doi.org/10.1016/j.neuroimage.2020.117074

Whitfield-Gabrieli, S., Nieto-Castanon, A., 2012. Conn: a functional connectivity toolbox for correlated and anticorrelated brain networks. Brain connectivity 2, 125–141. https://doi.org/10.1089/brain.2012.0073

Whitney, C., Kirk, M., O’Sullivan, J., Lambon Ralph, M.A., Jefferies, E., 2011. The neural organization of semantic control: TMS evidence for a distributed network in left inferior frontal and posterior middle temporal gyrus. Cerebral Cortex 21, 1066–1075. https://doi.org/10.1093/cercor/bhq180

Whitney, C., Kirk, M., O’Sullivan, J., Ralph, M.A.L., Jefferies, E., Lambon Ralph, M.A., Jefferies, E., 2012. Executive Semantic Processing Is Underpinned by a Large-scale Neural Network: Revealing the Contribution of Left Prefrontal, Posterior Temporal, and Parietal Cortex to Controlled Retrieval and Selection Using TMS. Journal of Cognitive Neuroscience 24, 133–147. https://doi.org/10.1162/jocn_a_00123

Wilson, M., 1988. MRC Psycholinguistic Database : Machine-usable dictionary, version 2 . 00. Behavior Research Methods, Instruments, & Computers 20, 6–10.

Yarkoni, T., Poldrack, R.A., Nichols, T.E., van Essen, D.C., Wager, T.D., 2011. Large-scale automated synthesis of human functional neuroimaging data. Nat. Methods 8, 665–670. https://doi.org/10.1038/nmeth.1635

Yeo, B.T.T., Krienen, F.M., Sepulcre, J., Sabuncu, M.R., Lashkari, D., Hollinshead, M., Roffman, J.L., Smoller, J.W., Zollei, L., Polimeni, J.R., Fischl, B., Liu, H., Buckner, R.L., 2011. The organization of the human cerebral cortex estimated by intrinsic functional connectivity. Journal of Neurophysiology 106, 1125–1165. https://doi.org/10.1152/jn.00338.2011.

Zhang, M., Wang, X., Varga, D., Krieger-Redwood, K., Margulies, D.S., Smallwood, J., Jefferies, E., 2020. Distinct default mode network subsystems show similarities and differences in the effect of task focus across reading and autobiographical memory. bioRxiv 2020.10.03.324947.

